# NKX2-5 congenital heart disease mutations show diverse loss and gain of epigenomic, biochemical and chromatin search functions underpinning pathogenicity

**DOI:** 10.1101/2025.06.20.659510

**Authors:** Alexander O. Ward, Nicole Schonrock, Alex J. McCann, Sabrina K. Phanor, Kian Hong Kock, Jesse V. Kurland, Fujian Wu, Nicholas J. Murray, James Walshe, Dimuthu Alankarage, Sally L Dunwoodie, Frederic A. Meunier, Mathias Francois, Martha L. Bulyk, Mirana Ramialison, Richard P. Harvey

## Abstract

Congenital heart defects (CHD) occur in ∼1% of live births, with inherited and acquired mutations and environmental factors contributing to causation. However, network perturbations in CHD remain poorly understood. We report an integrated functional-epigenomics approach to CHD, focusing on the cardiac homeodomain (HD) transcription factor NKX2-5, mutations which cause diverse heart structural and conduction defects. We selected twelve NKX2-5 CHD variants affecting different residue classes - homeodomain DNA base-contacting, backbone-contacting and helix-stabilizing, and those affecting other conserved protein:protein interaction (PPI) domains. In HL-1 cardiomyocytes, we profiled genome-wide DNA targets of NKX2-5 wild type (WT) and variants, their DNA binding affinity and specificity, PPI with known NKX2-5 cofactors and chromatin search dynamics. Variants showed diverse yet class-specific behaviours. All variants failed to bind many WT targets but retained binding to a subset of core cardiomyocyte-related targets, and bound hundreds of unique “off-targets” via changes to DNA binding site specificity, homodimerization, cofactor interactions and chromatin search functions. Our data suggest that complex residue-by-residue scale epigenomic, biochemical and chromatin search defects involving both loss-and gain-of-function contribute to CHD. These findings may inform precision molecular therapeutic approaches in patients with CHD.

## INTRODUCTION

Congenital heart disease (CHD) is the most common congenital abnormality in humans, occurring in ∼0.5-1% of all live births. Causative variants for CHD have been identified in a diverse set of genes known to coordinate cardiac development, including those for cardiac transcription factors (TFs) NKX2-5, TBX5, TBX20 and GATA4 ^1–5^, intercellular signalling intermediates, ciliary proteins ^6–10^, epigenetic regulators ^11–17^ and structural proteins ^18^. However, the precise mechanisms of causation for CHD remain poorly understood. TFs are of particular interest because they are conserved spatiotemporal drivers of gene regulatory networks (GRNs) ^19^ and interact dynamically with enhancers through an embedded motif grammar hard-wired into the genome ^20^.

Core cardiac TFs interact directly with each other as well as with a host of cofactors, whose genes are recursively wired into the network ^19,25–29^. Mutations in cardiac developmental TFs lead to catastrophic effects on heart development. For example, *Nkx2-5* null embryos show lethal arrest of heart development at early heart looping stages due to failures of cardiac field, chamber, vessel and conduction system differentiation and proliferation ^30–36^. There is no obvious genotype-phenotype correlation for NKX2-5 variants ^37^, similar to CHD-causing variants in other cardiac TFs such as *TBX5* and *GATA4*, which show overlapping phenotypic spectrums ^39,40^.

Most variants in cardiac TFs lie in conserved functional domains and are predicted to confer loss-of-function (LOF) effects via haploinsufficiency. The partial penetrance and expressivity of variants suggest complex influences on CHD outcomes, including dominant-negative actions, gains-of-function, modifier genes ^11,12,42,43^ and increased stochasticity of network outputs when TFs act close to functional thresholds ^41,44^. In true LOF, variant proteins would be unstable or incapable of performing their normal function. However, recent evidence suggests gain-of-function (GOF) effects for dominant TF variants in normal and cancerous cells ^25,29,45–52^, among them being changes to DNA binding specificity ^53^. We have previously shown that one NKX2-5 CHD-causing variant (Y191C) and a synthetic variant completely lacking the homeodomain (HD) retain binding to a subset of normal NKX2-5 targets and also bind many unique “off-targets” leading to their modified expression ^25^. Based on these findings, we hypothesise that dominant-acting TF variants act via LOF hand-in-hand with pervasive GOF, to disrupt and destabilise GRNs ^25^.

We report here a broader structure-function analysis of NKX2-5 CHD-causing variants identified through clinical genetics testing (Figure 1A). We selected 12 variants affecting different functional classes of residue within the NKX2-5 HD and other conserved PPI domains, and determined their target genes, DNA binding affinities and specificities, cofactor interactions and chromatin space search behaviours. Whereas variants showed individual properties, sub-classes were identified that targeted the genome in similar ways. All variants showed GOF effects, in some cases through a clear regulatory logic. Their unique genomic signatures reveal how different variant sub-classes perturb the cardiac GRN in different ways at single-residue scale, likely contributing to clinical manifestation of CHD.

**Figure 1.**
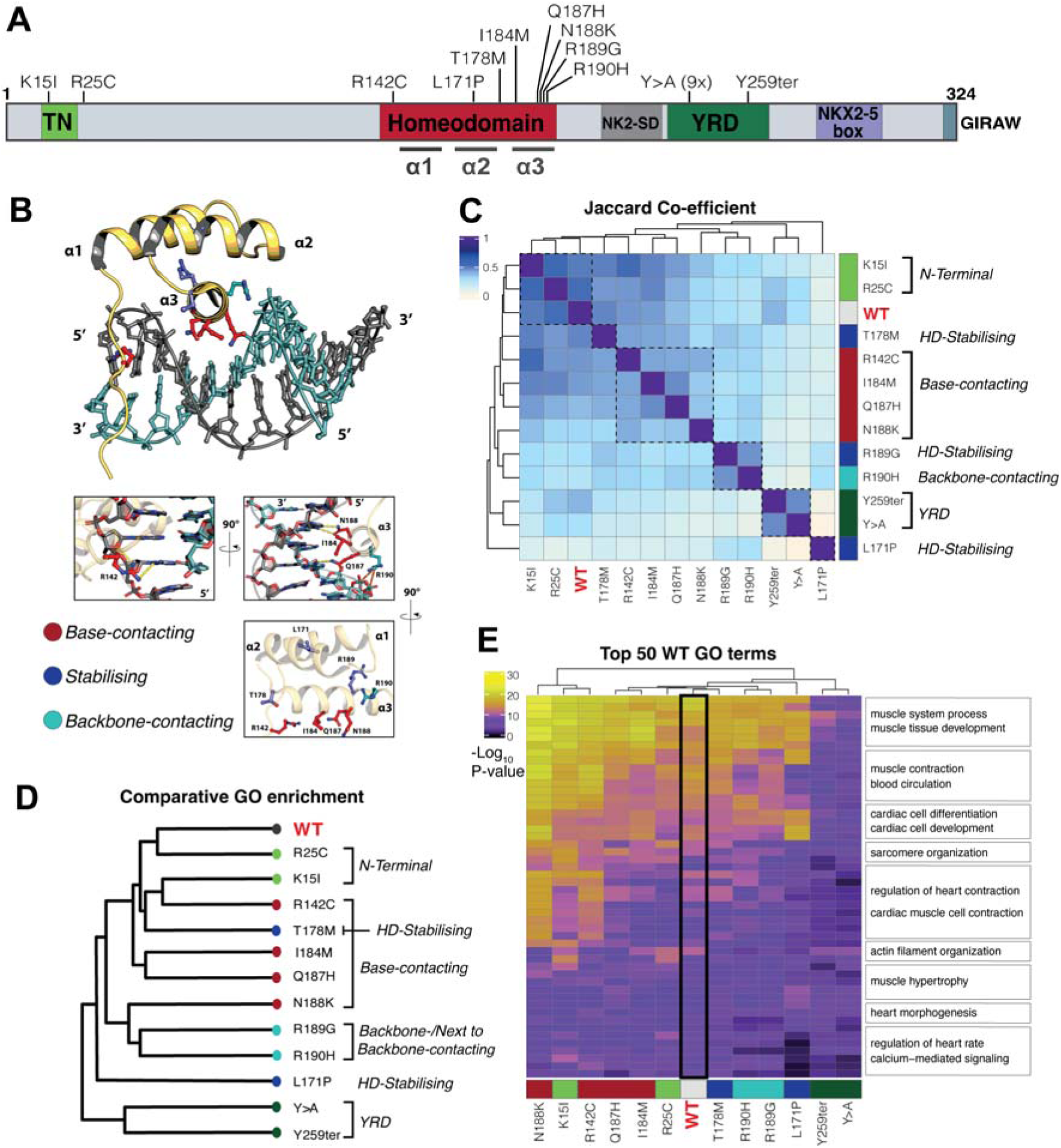
DamID reveals that Nkx2-5 allelic variants affect genomic targeting behaviours in a residue function specific manner (A) Schematic showing the amino acid positions of Nkx2-5 allelic variants. (B) Crystal structure of the Nkx2-5 WT Homeodomain (HD) with mutated residues highlighted by their functional class. (C) Degree of peak overlap of all WT and variant peaks measured using the Jaccard similarity co-efficient (between 0-1), with functional classes indicated and colour coded. (D) Unsupervised clustering of Nkx2-5 variants by the statistical similarity and log Odds Ratio scores of gene ontology (GO) terms, calculated using CompGO. (E) Significance values of the top 50 slimmed WT GO terms and corresponding significance for allelic variant, black indicates not significantly enriched.

## MATERIALS AND METHODS

### Variant scoring with CADD, BayesDel and REVEL

Using the list of 143 pathogenic *NKX2-5* variants collated previously ^37^, a VCF file was generated and uploaded to the Combined Annotation Dependent Depletion (CADD) online platform (version 1.6) ^55^. The Raw CADD score was extracted and plotted against nucleotide variant position using custom R scripts. We also used the BayesDel and Rare Variant Ensemble Learner (REVEL) as implemented in Ensemble Variant Effect Predictor (VEP) ^56,57^ to assess pathogenicity of NKX2-5 variants.

### Plasmids and Cloning

Sequences coding for cardiac TFs *Nkx2-5, Gata4, Tbx5, Tbx20* and *Hand1* were amplified from HL-1 cell cDNA, as described ^25^. *Nkx2-5* and other cardiac TF cDNA were cloned into pCR8/GW/TOPO vectors with the pCR/GW/TOPO TA Cloning kit (Thermo Fisher, MA, USA). Mouse *Nkx2-5* variant sequences were generated using site-directed mutagenesis of *Nkx2-5* cDNA and Gateway-cloned into DamID lentiviral vectors pLgw-EcoDam-V5 ^58,59^, a gift from Bas van Steensel (Addgene plasmid #59210). Vectors used for lentivirus production were obtained from Prof Didier Trono: pMLDg/pRRE (Addgene #12251), pRSV-Rev (Addgene #12253), and pMD2.G (Addgene #12259)). For Yeast-2-Hybrid assays, *Nkx2-5* WT or variant cDNA was cloned from pCR8/GW/TOPO vectors into pGADT7 AD library vectors and *Nkx2-5* WT or other cardiac TF cDNA (listed above) were cloned into pGBKT7 DBD bait vectors (Clontech, Mountain View, USA). *Nkx2-5* WT or variant cDNA was Gateway-cloned into pDEST15-GST bacterial expression vectors, kindly gifted by Robert Shearer (Garvan Institute for Medical Research), for production of proteins used in protein binding microarray (PBM) experiments. Lastly, for single molecule tracking experiments, *Nkx2-5* WT or variant cDNA was cloned in frame with HaloTag (amplified from pENTR4-HaloTag (w876-1); Addgene plasmid #29644; gifted by Dr Eric Campeau), into the CMV-promoter driven vector, pCMX-GW1, obtained from Dr Gavin Chapman (Victor Chang Cardiac Research Institute), using the NEBuilder HiFi DNA assembly kit (New England Biolabs (NEB), MA, USA).

### Cell Culture

The HL-1 cell-line were donated by Prof W C Claycomb (Department of Biochemistry and Molecular Biology, Louisiana State University, New Orleans, LA, USA) ^60^. HL-1 cells were maintained in Claycomb Medium (Sigma-Aldrich, MO, USA; #51800C) supplemented with 0.1mmol/L norepinephrine, 100μg/mL Penicillin/Streptomycin, 2mmol/L L-Glutamine and 10% Foetal Bovine Serum. HEK293T, HeLa and Cos-7 cells were acquired from ATCC (Rockville, MD) and cultured in Dulbecco’s Modified Eagle Medium (DMEM) supplemented with 100μg/mL Penicillin/Streptomycin, 1% GlutaMAX (all Gibco) and 10% Foetal Bovine Serum (FBS) (Sigma Aldrich), at 37°C with 5% CO_2_. Alterations to culture conditions will be detailed below.

### Liquid Yeast-2-Hybrid assay

Yeast-2-Hybrid assays were performed in liquid cultures as described ^25^. 1 μg pGADT7-AD-Nkx2-5 (WT or variant), containing the Gal4 activation domain, and 1μg pGBKT7-DBD-TF (NKX2-5, GATA4, TBX5, TBX20b or HAND1), containing the Gal4 DNA-binding domain, were co-transformed into chemically competent *S. Cerevisiae* (Clontech), using 240 μL PEG 3350 50%, 36 μL Li-Acetate 1.0 M, 25 μL Boiled SS-carrier DNA, 54 μL water. Double transformed cells were selected for growth on low stringency (-Leu/-Trp) selection plates at 30°C for 3 days, colonies were then picked and cultured in low-stringency liquid medium for a further 3 days at 30°C. Cultures were then selected for interactions in high stringency liquid medium (-Ade/-His/-Leu/Trp) at a 1:10 ratio of low stringency culture:high stringency medium and optical density (OD) of cultures was normalised. Fluoroscein Di-β-D-Galactopyranoside (FDG) was added to high-stringency medium to allow accurate fluorometric recordings of culture growth. Fluorescence was recorded at two time-points after after high stringency cultures (e.g. day 2 and day 3). Some interactions (NKX2-5 with HAND1 and GATA4) persisted beyond day 3 and were maintained until later time-points. Fluorescence recordings from time-point two (T_2_) were normalised to time-point one (T_1_).

### DNA-adenine methyltransferase identification (DamID)

DamID experiments were performed with modification of published protocols ^59^ as described ^25^. Briefly, confluent HL-1 cells were transduced with lentiviral vectors allowing undetectable expression of Dam-Nkx2-5 (WT or variant) fusion proteins driven by an uninduced *heat shock protein 68* promoter. After 40 hours, genomic DNA was extracted using a Gentra PureGene Cell kit (QIAGEN, Venlo, Netherlands), digested by *Dpn*I at 37°C for 6 hr, and amplified by ligation-mediated PCR. PCR products were further fragmented with DNase I at 24°C for 1 min, to lengths of between 200-2000 bps, and labelled and hybridised to Affymetrix mouse 1.0R promoter microarrays according to manufacturers’ instructions. Three independent DamID experiments were performed utilising biological triplicates.

### DamID bioinformatic analysis

#### Array processing, peak detection and gene assignment

Array processing, peak detection and gene assignment were performed as previously described in Bouveret *et al*., 2015 ^25^. Briefly, quality control was conducted using the ‘affy’ and ‘affyPLM’ R packages to assess probe-level distributions via RLE and NUSE plots, with outlier arrays excluded. Probe remapping to the mm9 genome (NCBI Build 37) was carried out using Starr, and array normalisation and peak calling were performed using CisGenome v2.0 with quantile normalisation and the TileMapv2 algorithm (minimum 8 probes, moving average ≥3.5). Peaks were assigned to genes using GREAT with “basal plus extension” settings (6.5 kb upstream, 2.5 kb downstream, max 100 kb), considering only non-coding regions covered by the array, with annotations retrieved from Ensembl API v66. Overlaps and distances to TSS were calculated using custom scripts and BEDTools.

#### Genomic footprinting

To evaluate similarity between transcription factor binding profiles across WT and variants, genomic footprinting was performed using the bedtools *bedr* R package (v1.0.7), which enabled efficient manipulation and comparison of genomic intervals in BED format. Peak files from each dataset were pre-processed to retain high-confidence regions within canonical chromosomes. Pairwise comparisons between all combinations of peak sets were conducted using the *jaccard* function in *bedr*, which computes the Jaccard index—defined as the size of the intersection divided by the size of the union of two sets of genomic intervals. This metric provides a scale-independent measure of similarity between binding profiles, capturing both shared and unique genomic occupancy. The resulting Jaccard indices were assembled into a symmetric similarity matrix representing all pairwise comparisons between samples or conditions. This matrix was visualised as a clustered heatmap using the *ComplexHeatmap* R package (v2.16.0) ^61^ which provides advanced layout and annotation capabilities for genomic data visualisation. Hierarchical clustering with complete linkage and Euclidean distance was applied to group samples according to shared binding patterns. Annotations denoting NKX2-5 variant allele were included to aid interpretation, allowing for identification of conserved, divergent, or condition-specific regulatory footprints across the dataset.

#### GO analysis

Gene ontology (GO) enrichment analysis was performed using the *clusterProfiler* R package (v4.8.1) to identify biological processes significantly associated with genes linked to binding regions. Gene identifiers were first mapped to Entrez IDs using *org.Mm.eg.db*, and enrichment analysis was conducted using the *enrichGO* function with the *Biological Process* ontology. The entire mouse genome was used as the background universe to provide an unbiased enrichment context. To aid interpretability, GO term redundancy was reduced using the *simplify* function in *clusterProfiler*, which collapses semantically similar GO terms while retaining the most statistically significant representative. Enriched terms with adjusted p-values ≤ 0.05 were considered significant.

To support interpretation, visualisation of enriched GO terms was performed using built-in plotting functions including dot plots, bar plots, and enrichment maps. Where appropriate, gene concept networks were also generated to highlight shared gene-level contributions to multiple GO terms. For comparative enrichment analysis across multiple datasets or conditions, the *CompGO* R package ^62^ was used. *CompGO* enables statistical comparison of GO term enrichment profiles by computing log odds ratios and associated confidence intervals between groups, facilitating direct assessment of whether specific biological processes are differentially enriched.

#### DNA motif discovery

Motif discovery and known motif enrichment analyses were performed using many of the same tools and procedures described in Bouveret *et al.*, 2015 ^25^. Briefly, Trawler_standalone ^63^ was used for de novo motif discovery, with default parameters except where noted. Motif discovery was performed on repeat-masked sequences ±6.5 kb from the transcription start site of genes associated with binding peaks. To reduce redundancy among highly similar motifs, Trawler output was further processed using the matrix clustering module of *RSAT* ^64^ which groups motifs into superfamilies based on similarity scores, allowing for a more interpretable set of representative motif classes.

To complement the original motif discovery, additional *de novo* motif analysis was performed using the *HOMER* suite (v4.11) ^65^ HOMER was run with default settings on peak-centred 200 bp sequences, and enrichment was calculated relative to a custom background set matched for GC content and length. *de novo* motifs identified by Trawler were further annotated using *TomTom* from the MEME Suite ^66^, which performs motif-motif comparisons against curated databases (e.g., JASPAR) to identify putative transcription factor matches. Matches with q-values < 0.05 were considered significant.

Known motif enrichment analysis was carried out using *Clover* as previously described, testing for overrepresentation of known transcription factor binding motifs in input peak sequences relative to a background composed of tiled regions from the Affymetrix Promoter 1.0R Array. Motifs with p-values < 0.05 were considered significantly enriched.

#### Network analysis

Protein–protein interaction networks were constructed using the STRING database (v12.0) ^67^, which integrates evidence from curated databases, experimental data, gene co-expression, and text mining to infer both direct (physical) and indirect (functional) associations. Gene sets derived from peak-associated loci or enriched GO terms were submitted to the STRING web interface, restricted to *Mus musculus*, using a minimum interaction confidence score of 0.7 (high confidence). STRING analysis was used to identify significantly enriched interaction networks, with enrichment p-values calculated relative to a genome-wide background. Functional clustering within the interaction networks was performed using STRING’s built-in k-means clustering algorithm, revealing coherent modules based on shared network topology and biological function. Networks were exported from STRING and visualised using *Cytoscape* (v3.10.0)^68^, enabling interactive exploration and refinement of network topology. Functional clusters identified by STRING were preserved and annotated within Cytoscape to highlight distinct biological modules. Node attributes such as gene function, expression level, or GO category were integrated into the visualisations to support interpretability and presentation.

#### Protein Binding Microarray (PBM) experiments

Glutathione S-transferase (GST)-NKX2-5 (WT or variant) fusions were expressed from bacterial expression vectors, driven by a T7 promoter, using the PURExpress *In Vitro* Protein Synthesis kit (NEB), as per the manufacturer’s instructions. Molar concentrations of GST-NKX2-5 fusion proteins were estimated using Western blotting and comparison against serial dilutions of recombinant GST (Sigma Aldrich, MO, USA). PBM experiments were performed as described^48^. Briefly, custom-designed, “all 10-mer” universal arrays in 8 x 60K, GSE format (Agilent Technologies; AMADID #030236) were double-stranded and used in PBM experiments with Alexa488-conjugated anti-GST antibody (Invitrogen A-11131) essentially as described previously ^69,70^. All proteins were assayed in at least duplicate at 200 nM final concentration in PBS-based binding and wash buffers. All three arrays analysed contained a Reference WT-NKX2-5 protein (WT1-3) to allow analysis against internal WT controls and cross-array comparisons. Specific analyses on the three arrays were as follows: v213: WT1, I184M, L171P, Q187H, R25C, R142C, T178M; v214: WT2, N188K, R189G, R190H, Y191C; v311, WT3, Y259ter. Arrays were scanned using a GenePix 4400A microarray scanner (Molecular Devices). At least 3 scans were taken for each slide at different photomultiplier tube (PMT) gain settings. Microarray data quantification, normalization, and motif derivation were performed essentially as described previously using the Universal PBM Analysis Suite and the Seed-and-Wobble motif-derivation algorithm ^69^.

#### Single Molecule Tracking (SMT) experiments

SMT experiments were carried out largely as described previously ^71^. Briefly, HeLa cells were seeded at a density of 1.5×10^5^ cells into 0.1% Gelatin-coated (Sigma Aldrich) glass-bottom 35mm FluoroDishes (World Precision Instruments, FL, USA) for 24 hours prior to transfection. Transfections were performed using XtremeGENE 9 transfection reagent (Roche, Switzerland), to introduce 1 μg plasmid DNA (pCMX-HaloTag-Nkx2-5 (WT or variant)), diluted in OptiMEM (Thermo Fisher) and complex was allowed to form, per manufacturer’s instructions. Transfections were incubated in FluoroBrite DMEM (referred to below as imaging medium) (Gibco), supplemented with 10% FBS and 1% GlutaMax for 24 hours at 37°C and under 5% CO_2_. Prior to imaging, cells were washed twice with Dulbecco’s PBS and labelled with 1nM JF549-HaloTag ligand (Promega, WI, USA) for 20 minutes in imaging medium at 37°C and under 5% CO_2_. The ligand was removed, cells were washed twice and fresh imaging medium was added. SMT images were acquired using an Elyra PALM/STORM (PS.1) super-resolution microscope with total internal reflection fluorescence (TIRF) architecture, using a 100x alpha Plan-Apochromat oil 1.46 NA objective and a BP 570-650/LP 750 filter cube (Zeiss, Germany). Samples were excited with an HR DPSS 531nm laser with a high-power TIRF filter and fluorescent signal was collected with the Andor iXon 897 EMCCD camera. Total power of the 531 nm laser line is 200 mW, at the low laser power used here, the power measured at the exit from the objective was 4.9 mW in epifluorescence mode, this value should be equivalent in oblique illumination mode. The oblique illumination angle used was 60.18°, equivalent to Highly Inclined and Laminated Optical sheet (HILO) illumination angle ^72^. Imaging with low laser power and HILO illumination ensured that cells did not show any signs of phototoxicity or significant photobleaching during imaging. Two modes of SMT imaging were performed using this setup. Firstly, fast SMT at 50 Hz (20ms framerate) was performed to acquire 6000 frames. This enables the tracking of single molecules with high temporal resolution (immobile and mobile populations). Secondly, slow SMT at 2 Hz (500ms framerate) was performed for 500 frames to remove background created by fast moving molecules and derive lifetime of the immobile population. Final image resolution for both modes was 0.1μm x 0.1μm x 1μm. Image frame sizes were reduced to capture only a single nucleus, without changes to pixel size, preventing the need to post-crop images. Prior to capture, each nucleus was pre-bleached for 10-30 seconds, to reduce background fluorescence and the density of HaloTag-fusion molecules.

### SMT data analysis: fast SMT (20ms)

#### SLIMfast tracking, SpotOn and diffusion coefficient calculation

Fast SMT image stacks were batch converted to tiff files using FIJI and analysed using similar pipelines to those published ^73,74^ and MATLAB version 2015a. The pipeline uses the MATLAB script *SLIMfast.m*, which employs a two-step process to localise molecules, through a modified version of the multiple-target tracing (MTT) algorithm^75^. Similarly, to McCann *et al*.^71^, batch processing was performed with an error rate of 10^-6.5^, a detection box of 7 pixels, maximum number of iterations of 50, a termination tolerance of 10^-2^, a maximum position refinement of 1.5 pixels, an N.A. of 1.46, a PSF scaling factor of 1.35, and 20.2 counts per photon, an emission of 590 nm, a lag time of 20 ms and a pixel size 100 nm. Trajectory creation was performed using the maximum expected diffusion coefficient of 1.5 μm^2^/sec, determined from SpotOn analysis (above) and qualitative assessment of trajectories (i.e. presence of no/very few incorrectly localised molecules). Trajectories with less than 5 tracks were excluded and cells with significant divergence from normal distribution were removed as outliers using interquartile range thresholding. SLIMfast tracked data was then processed with custom MATLAB scripts to output log_10_ diffusion coefficient measurements and trajectory maps from ROIs of each cell (whole nucleus). Diffusion coefficient measurements for all cells were combined with another custom MATLAB script and plotted in GraphPad Prism.

#### TrackIt tracking and, kinetic modelling

Fast SMT image stacks were loaded directly into TrackIt v1.1 ^76^ using MATLAB version 2020b. TrackIt provides a fully-integrated interface for visualisation, processing and analysis of SMT ^76^. Regions of interest (ROIs) were drawn around each nucleus to remove cytoplasmic and extracellular background signal. The nearest neighbour tracking algorithm was used with a threshold of 1.5, a tracking radius of 9 pixels, a minimum trajectory length of 5 tracks and a maximum of 2 gap frames between trajectories. Mean Squared Displacement (MSD) was fit with a power law function. 90% of tracked points and an offset of 0.5 were used for MSD fitting. Cells with significant divergence from normal distribution were removed as outliers using interquartile range thresholding. Fitted data was exported for principal component analysis (PCA) and dwell time survival function plotting, using custom R scripts. Raw tracked data for each cell was output for use in downstream analyses detailed below. Firstly, kinetic modelling was used to profile the diffusion coefficients of specific populations of molecules, and the fraction of total molecules in each population, using the SpotOn MATLAB package, developed by Hansen and colleagues ^77^, run using MATLAB version 2020b. We used the 3-state/population model, which categorises molecules into: Bound, Slow-diffusing or Fast-diffusing states, which allows the capturing of complex nuclear dynamics seen in many TFs. TrackIt tracked output files for individual cells were used in SpotOn, and a 3-state model was fit with default settings, except for the following changes: a framerate of 20ms, 6 timepoint delays considered, entire trajectories were used for fitting, 2 gap frames were allowed, 10 fitting iterations and the localisation error was fit from the data. In addition, the thresholds for min/max diffusion constants (in μm^2^/sec) were constricted as follows, to prevent incorrect fitting: Bound = 0.00001-0.1; Slow = 0.001-0.5; Fast = 0.2-5. Population data were then collated and plotted using custom R scripts.

#### SMT data analysis: slow SMT (500ms)

Slow SMT image stacks were batch converted to tiff files using FIJI and analysed using similar pipelines to those published ^78^, run with MATLAB version 2015a. The pipeline uses the MATLAB script *SLIMfast.m*, which employs a two-step process to localise molecules, through a modified version of the multiple-target tracing (MTT) algorithm^75^. SLIMfast batch processing was performed with an error rate of 10^-6.5^, a detection box of 7 pixels, maximum number of iterations of 50, a termination tolerance of 10^-2^, a maximum position refinement of 1.5 pixels, an N.A. of 1.46, a PSF scaling factor of 1.35, and 20.2 counts per photon, an emission of 590 nm, a lag time of 500 ms and a pixel size 100 nm. Trajectory creation was performed using the maximum expected diffusion coefficient of 0.05 μm^2^/sec, determined from SpotOn analysis (above) and qualitative assessment of trajectories (i.e. presence of no/very few incorrectly localised molecules). Following this, a modified version of the *Calculate length_2fitting_v3.m* MATLAB script was used, taken from McCann and colleagues ^71^, with script provided by Dr Zhe Liu. Briefly, the modified script batch calculates the lifetime of molecules for each cell and fits a one-and two-component exponential decay function, outputting decay curves. These decay curves provide the average dwell time for specific and non-specific fractions, and the ratio between both fractions. Cells with significant divergence from normal distribution following slow tracking analysis were removed as outliers using interquartile range thresholding. Dwell times for specific and non-specific fractions, and the ratio, were output directly and collated and plotted in GraphPad Prism.

## RESULTS

### Mutational landscape of NKX2-5

NKX2-5 CHD variants are spread across known conserved domains – the N-terminal trans-repressive *Tinman* (TN) domain, DNA-binding homeodomain (HD), and C-terminal NK2-Specific Domain (NK2SD), Tyrosine Rich Domain (YRD) and NKX2-5 Box^79,80^ (Figure 1A). We selected 143 natural variants previously associated with diverse CHD clinical phenotypes ^37^ and assessed their predicted pathogenicity *in silico* using Combined Annotation Dependent Depletion (CADD) ^81^ (Figure S1A). All HD variants were predicted to be highly pathogenic. However, missense variants within specific HD helices (H1-H3) and those lying between helices (inter-HD) varied considerably with respect to clinical phenotype and CADD scores; thus, HD helical position carries distinct functional outputs ^53^. Barrera *et al*. ^48^ previously showed that HD residues directly contacting DNA bases or backbone phosphates, or situated adjacent to these residues, were significantly enriched among Mendelian disease mutations. Therefore, for increased granularity, we classified HD variants according to their predicted HD residue functions as DNA-base-contacting, DNA-backbone-contacting or HD-stabilising, rather than their overall helical position ^48^ using the NKX2-5 HD crystal structure as reference (Figure 1B)82.

A list of variants was chosen for in-depth profiling based on: 1) association with CHD clinical phenotypes; 2) classified as *Pathogenic* or *Likely Pathogenic* by the American College of Medical Genetics and Genomics (ACMG) ^83^; 3) affecting different sub-classes of HD residue (DNA base-contacting, DNA backbone-contacting, HD stabilising); 4) inclusive of a broad range of predicted pathogenicity scores; and 5) previously investigated *in vitro* (Figure 1A). We include natural variant K15I located in the TN trans-repressor domain as a ‘mild’ variant ^84^, and one previously characterised severe “synthetic” variant impacting the conserved YRD located downstream of the HD (Figure 1A), in which we converted all nine conserved tyrosine residues of the tyrosine-rich YRD to alanine (Y>A)^79^, complementing a natural stop-gain clinically-relevant variant in the YRD (Y259ter). Clinical associations for the eleven natural variants are shown in Figure S1B. Pathogenicity predictions using REVEL ^57^ and BayesDel ^56^ showed a similar spectrum of scores to CADD (Figure S1C). All variants localised to the nucleus in transfected Cos-7 cells, which lack endogenous NKX2-5 expression (Figure S2).

### Genome-wide target signature of NKX2-5 variants is linked to functional class of mutation

Profiling of genome-wide targets of NKX2-5 WT and variants was performed using DNA-adenine methyltransferase identification (DamID) (Figure S3A) ^59,85–87^ which avoids the need for high quality antibodies and artefacts due to chromatin crosslinking ^88^. NKX2-5 WT or variant allele N-terminal fusions with bacterial DAM methylase were expressed at trace levels in the HL-1 cell line (derived from atrial cardiomyocytes), as reported ^25,89,90^. Targets were detected by hybridizing GATC-me DamID target fragments to Affymetrix gene promoter microarrays covering ∼10 kb upstream and downstream of the transcription start site (TSS), in triplicate. We identified 1573 NKX2-5 WT peaks with low false discovery rates (FDR) (<0.05 cutoff; Figure S3B-C). Around half of NKX2-5 variants led to a significant reduction in peak numbers (Figure S3B). Despite significant loss of targets, the distributions of WT and variant targets across genome features were broadly similar (mostly within 5 kb of the TSS), suggesting broad retention of binding to genome regulatory elements (Figure S3C-D).

We first compared the peak overlap of WT and variants using the Jaccard coefficient (JC), which measures the similarity of peaks by calculating the ratio of the number of intersecting base-pairs to the total number of base-pairs within peak sets, minus the intersection ^91^. This statistical method revealed that peak footprints of variants within the same functional class overlapped well with one another (higher JC) (Figure 1C). For example, the two NKX2-5 variants located at the N-terminus of the protein - K15I and R25C (light green; Figure 1C) - showed very similar peak footprints to one another and to NKX2-5 WT. In contrast, both naturally occurring Y259ter and synthetic Y>A variants in the YRD, a known protein-protein interaction (PPI) domain (dark green) ^25^, showed highly similar peak footprints, that were distinct from WT and all other variants (Figure 1C).

The genomic footprints of HD variants were much more nuanced, likely reflecting the underlying complexity of structure-function relationships across HD residues (Figure 1C) ^25,29,48,74^. Most DNA base-contacting residue variants clustered together, albeit showing a graded divergence from WT (Figure 1C, red) with N188K the most severe affected. Notably, R190H, the single DNA backbone-contacting variant chosen, was more severely affected than *all* DNA base-contacting variants. R190H clustered together with its adjacent variant R189G at a HD-stabilising position, and both were highly distinct from WT and other HD stabilising variants. As they appear functionally similar across virtually all analyses in this study, for simplicity we assigned both R190H and R189G as “backbone-contacting” (Figure 1C). Two other variants at HD-stabilising positions, T178M and L171P, showed highly divergent impacts, with T178M closer to WT and L171P (H2 helix-breaking ^92^) the most distal to WT of all variants profiled.

Following assignment of unique genes to each DNA binding peak detected using GREAT ^93^, WT and variant gene-sets were compared using the Spearman correlation coefficient. The resulting map showed highly similar groupings to that observed using the Jaccard score, especially for more severe variants (Figure S4A). Hierarchical clustering and principal component analysis (PCA) of Gene Ontology (GO) terms associated with WT and variant gene-sets also demonstrated the granularity between variant subclasses (Figure 1D; Figure S4A-B). As a further test, we visualised how cardiac muscle-related GO terms associated with variant sub-class genes compared to those of NKX2-5 WT (Figure 1E). Three variants, namely the N-terminal variant K15I, and two base-contacting mutations N188K and R142C, were associated with gain of cardiac muscle GO term significance, suggesting a gain-of-function effect. In contrast, R190H and R189G backbone-contacting variants, helix-breaking L171P variant and both YRD variants were associated with a broad reduction of muscle GO term significance and a reduction in overall number of target peaks. Collectively, these data highlight the profoundly different types of structure-function relationship within NKX2-5, some distinguished at single residue resolution.

### NKX2-5 variants disrupt gene-regulatory logic in a functional domain-specific manner

Using Trawler ^63,94^, a total of 175 unique *de novo* motifs were predicted for NKX2-5 WT and all variant peaks. We clustered these motifs semantically using RSAT ^64,94,95^, identifying 10 distinct sequence-related clusters (#1-10; Figure 2A), then compared these against known motifs using Tomtom ^66^. Variations of the high affinity NKX2-5 binding motif (NKE) were prominent (cluster 1; Figure 2A,B). The Nuclear Factor 1 (NF-1) motif was the dominant feature in cluster 2, more prevalently than the NKE itself, consistent with our previous findings ^25^. Cluster 3 represented a NK2 HD half site. Motifs for other TF family members occurred less frequently. To visualise graphically, we compared motif enrichment significance levels (Z-scores) in WT and variant peaks for all motif clusters (Figure 2C). In peaks for TN domain variants K15I and R25C, there was highly significant over-representation of the high affinity NKE (dominant in cluster 1) and NF-1 motifs (NF-1X cluster 2), in common with NKX2-5 WT peaks, consistent with their similar DamID signatures. For HD mutations, the NKE enrichment score varied considerably, with R142C (base-contacting) similar to WT, and severe mutations N188K and L171P showing no significant enrichment (Figure 2C). The YRD variants also showed a low NKE enrichment score despite carrying an intact HD. Generally, there was concordance of enrichment for the NF-1 motif and NKX2-5 motifs across variants, suggesting a close functional relationship between NKX2-5 and NF-1 factors on chromatin, aligned with our previous finding that most NF-1 isoforms are interacting cofactors of NKX2-5 ^25^. N188K and R189G showed strong Z-scores for NF-1 motifs and lower Z-scores for NKE-related motifs, suggesting that for some ofthese target peaks NKX2-5 binds indirectly. The YRD variant showed low Z-scores for both NKE and NF-1 motifs, suggesting that the YRD is necessary for interaction of NKX2-5 with NF-1 proteins. Cardiac TF family motifs MEF2 and HAND were significantly enriched in certain variant target sets as a gain-of-function, e.g., the MEF2C motif showed highest Z-scores for base-contacting residue variants I184M and Q187H, which have similar target gene profiles (Figure 1C,D).

**Figure 2.**
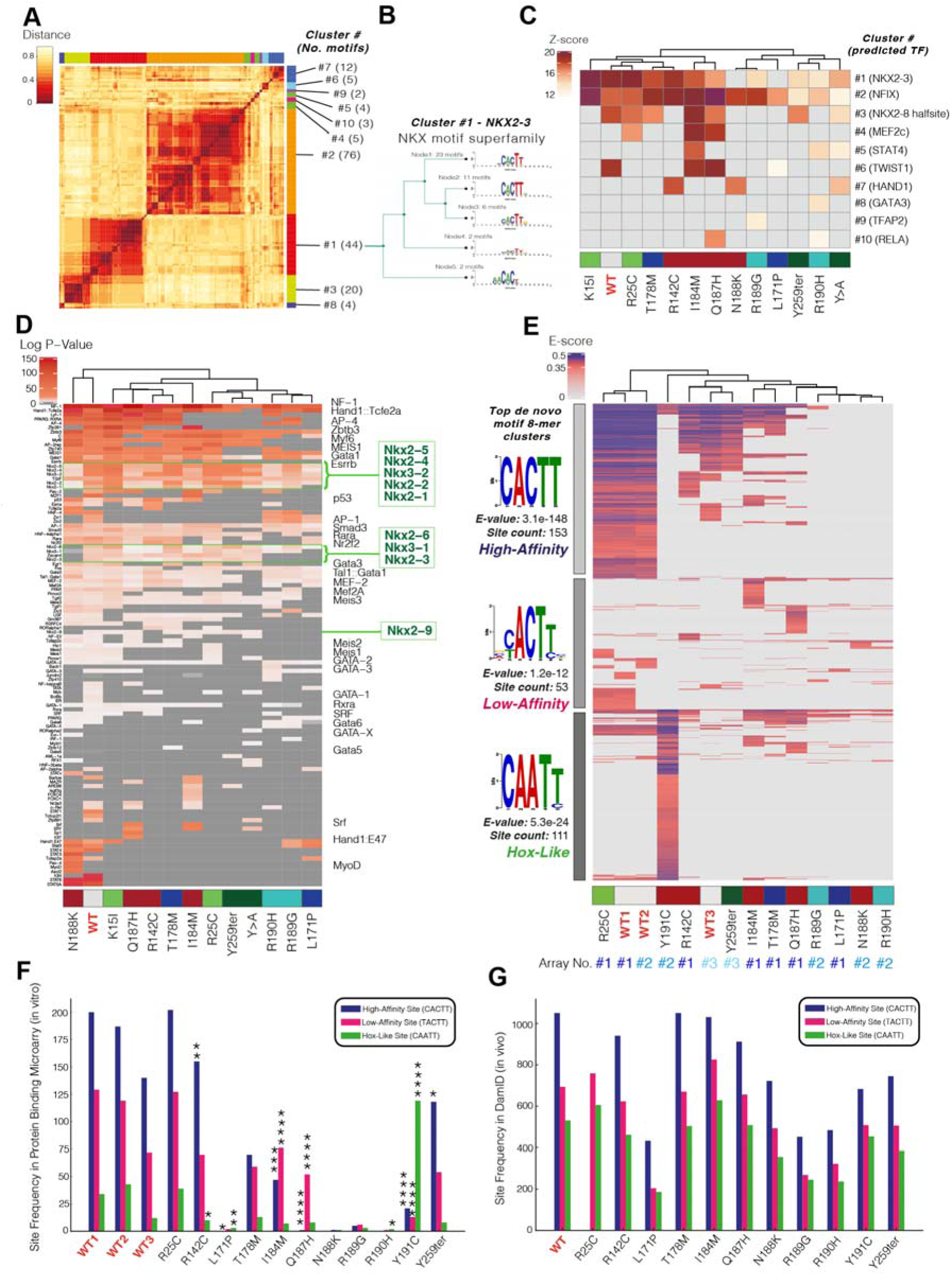
Disruption of DNA binding occurs in a residue function specific manner in Nkx2-5 variants (A) Distance matrix for all de novo discovered motifs identified using *Trawler* (values between 0-1) and motif clusters identified with RSAT matrix clustering. (B) Example of a reduced dendrogram for cluster #1, the NK2-type motif, highlighting the number of individual motifs discovered in each node (groups of similar motifs) and aggregate position weight matrix (PWM) for each node. (C) Significance (Z-score) of each de novo motif cluster identified with *Trawler*. (D) Significance (log p-value) of each known motif identified with *Clover*, with all motif names present on left and only cardiac-related motifs present on the right. (E) E-score distribution of all significantly bound 8-mers (above cut-off of 0.35) in 3 independent protein binding microarrays (PBM), with 3 major hierarchical clusters of motifs identified on the left (High-affinity, Low-affinity and Hox-like). (F) Frequency of three major detected 5-mer motifs (High-affinity, Low-affinity and Hox-like) in *in vitro* PBM 8-mers, across WT and all variants. (G) Frequency of three major detected 5-mer motifs (High-affinity, Low-affinity and Hox-like) in *in vivo* DamID peaks, across WT and all variants. Significance in (F and G) was calculated with a Two-sided Fisher’s exact test, * indicates p<0.05, ** indicates p<0.01, *** indicates p<0.001 and **** indicates p<0.0001 (not significant not indicated).

We used Clover ^96^ to directly assess known motif enrichment, finding strong enrichment of cardiac TF motifs (GATA, MEIS, HAND, MEF2, MYOD (bHLH), MYF6 (bHLH), and SRF) for NKX2-5 WT targets, as expected (Figure 2D). Again, we observed enrichment of NF-1 and NKX motifs across variants to different degrees. Other cardiac and muscle-related TF motifs and those for auxiliary co-factors (RARA, RXRA, AP-1, and p53) were also notably enriched, albeit in mosaic patterns.

### Analysis of NKX2-5 WT and variant DNA binding in vitro and in vivo

We next investigated DNA-binding affinity and specificity of NKX2-5 WT and a selection of variants using *in vitro* protein binding microarrays (PBM) incubated with full-length NKX2-5-glutathione S-transferase (GST) fusion proteins ^69,97,98^. PBM allows comprehensive analysis of the *in vitro* DNA binding preference of TFs among all possible DNA sequences of specified lengths. DNA binding affinity is reflected by enrichment (E)-scores, which are rank-based and therefore robust to comparisons between PBM chips of different design, scale and sensitivity ^69,70^ (note that one PBM chips, comparing NKX2-5 WT with Y259ter, was less sensitive than others; Figure 2E). Initial analysis of position weight matrices (PWM) identified a palindromic motif composed of two NKX half sites (AAGT/ACTT) as the primary 8-mer across three experiments, and the core NKE motif (CACTT) as the secondary 8-mer (Figure S5A-B). However, the core NKE was bound more significantly by WT NKX2-5 (higher E-score) (Figure S5C-D), and the palindrome was not detected in DamID targets. All variants except for R25C showed a reduction in binding to the NKE (Figure S5D). The pattern of binding to the NKE and half site palindrome were similar, although Q187H (backbone-contacting) had higher affinity for the palindrome.

We next compared the most significantly bound 8-mer classes (E-score >0.35; false discovery rate <0.01 ^70^) between WT and variants (Figure 2E). Almost all variants, including those affecting residues outside of the HD, showed reduced binding affinity (fewer bound 8-mers, lower E-score), or altered specificity (deviation from the WT 8-mer binding patterns). Hierarchical clustering segregated three major 8-mer classes (grey bars in Figure 2E), which corresponded to the high-affinity NKE (classical NKX2-5 consensus binding sequence), low-affinity NKE (variation of the consensus sequence), and HOX-like homeodomain TFBS clusters. NKX2-5 binding to the high-affinity consensus NKE sites and HOX-like sites with lower affinity was shown previously ^97,99^. Different variants displayed unique binding patterns with all except R25C)showing broad loss of binding across all three 8-mer classes, but also gain of binding to non-WT 8-mers (Figure 2E). For base-contacting variant Y191C, 8-mers were strikingly enriched in HOX-like sites, consistent with our previous DamID findings ^25^. This is because Y191 is essential for the ability of helix 3 of NK2-class HDs to extend by 20 amino acids, allowing them to adopt an ordered state and bind at high affinity to the NKE ^100^. Base-contacting variants Q187H and I184M and HD-stabilising residue variant T178M showed reduced binding to high-affinity NKE sites and increased binding to unique low-affinity NKE sites and HOX-like sites. Proportional analysis of binding site distribution in PBM (Figure 2F) validated the significance of Y191C binding to HOX-like sites and a shift in binding preference for I184M and Q187H from high-to low-affinity sites. Strikingly, R142C and Y259ter demonstrated the opposite trend, showing proportionally reduced binding to low-affinity NKE and HOX-like sites and a preference for high-affinity NKE sites in comparison to WT.

We next assessed whether a comparable shift in binding site distribution was evident *in vivo*. All three classes of motifs were found within NKX2-5 WT and variant DamID target peaks (Figure 2G). However, the proportionality changes found by PBMs were not seen in DamID peaks, suggesting that the observed affinity/specificity shifts observed *in vitro* are compensated for *in vivo* by supporting co-factor interactions and chromatin structure. Such compensation may contribute a degree of robustness to the cardiac GRN in the face of genetic perturbations.

### Influence of motif flanking 5’ and 3’ bases on DNA binding affinity in vitro and in vivo

We next explored the effects of 5’ and 3’ bases flanking the NKE core site (CACTT) on regulatory logic. In PBM experiments (Figure S5E), WT NKX2-5 showed a clear preference for a 5’-C and 3’-A flanking bases, whereas a 3’-T and other flanking base conformations reduced binding affinity (Figure S5F-M). To investigate how NKE flanking base preference impacted the cardiac GRN *in vivo*, we selected NKX2-5 WT DamID peaks for the occurrence of high and low affinity sites (Figure S5N-P). GO term analysis of associated genes revealed a significant functional segregation of high-(**<u>C</u>**CACTT) and low-affinity flanking (CACTT**<u>T</u>**) sites (Figure S5N,O). Genes close to high affinity sites were highly enriched in GO terms for cardiomyocyte developmental, structural and functional attributes (Figure S5N). Targets close to low-affinity sites did not show strong muscle signatures and segregated with more peripheral dimensions of the cardiac GRN, such as metabolism and extracellular matrix, albeit non-significantly (Figure S5O). Most NKX2-5 variant peaks were more significantly enriched in low-affinity flanking sites (Figure S5P), with exceptions being R25C and T178M, consistent with these variants having more WT-like target and GO term profiles (Figure 1C-E). Thus, core-flanking bases exert a significant influence on NKX2-5 WT binding affinity and cardiac GRN logic, as has been found for other TFs ^101–104^. Furthermore, the binding signatures of most of the profiled NKX2-5 variants, including those outside the HD, show a preference for lower affinity sites, likely influenced by changes in intrinsic DNA binding preferences, and other specificity factors, such as co-factor interactions.

### Variants within the YRD fail to bind cardiac regulatory genes due to loss of co-factor interactions

We next explored in detail the structure-function consequences of NKX2-5 point mutations. We focused initially on the two variants within the YRD that showed severe network consequences. Y259ter is a stop-gain variant linked to multiple CHD phenotypes in humans, including atrial and ventricular septal defects, and double outlet right ventricle ^37^. Y>A is a synthetic variant originally designed to probe the function of the YRD in mice, whereby all tyrosines of the YRD were converted to alanine ^79^. We first used CompGO, a tool for comparative GO term enrichment combined with the Jaccard coefficient and correlation analysis to statistically compare GO term enrichments between genes associated with WT and all YRD variant peaks ^62^ (Figure S7A-B). Results indicated that YRD variant GO terms diverged from those of WT, with Y259ter.

We categorised WT and variant peaks for YRD variants into three subsets based on their overlap, as previously described ^25^ (Figure 3A-B). A-sets were exclusively bound by WT but not variants; B-sets were bound by both WT and variants (retained targets); C-sets were uniquely bound by variants, which we previously termed “off-targets”. Comparing replicate experiments for NKX2-5 WT performed 2 years apart (n=3-4), we have compellingly demonstrated that A sets (lost targets) and C sets (off-targets) are not due to experimental noise ^26^. A-set target genes for YRD variants showed over-representation of GO terms for biological processes related to *striated muscle development* and *differentiation* and a high degree of overlap (70% and 85% for Y>A and Y259ter, respectively) (Figure 3C-E; Figure S7C,D), highlighting that YRD variants, even though retaining an intact DNA binding domain (HD), fail to bind elements of the core cardiac GRN. Interaction network analysis at the gene level using STRING ^105^ revealed that YRD variant A-set genes displayed networks with high significance/connectivity (Table S2). Overlapping lost (A-set) genes between Y>A and Y259ter also had a highly significant STRING PPI networks enrichment p-value of 6.54e-13. Network analysis using Markov clustering ^106^ revealed that cardiac-related genes, such as *Scn5a, Tbx5* and *Ryr2*, were at the core of a larger network (Figure 3E). Other key network clusters in YRD lost targets included genes related to *extracellular matrix* and *insulin signalling* (*Irs1, Itga7, Timp3* and *Tgfb3*), as well as those for *protein removal* and *apoptosis* (*Ubc, Ube1, Nfkb1* and *Bcl10*).

**Figure 3.**
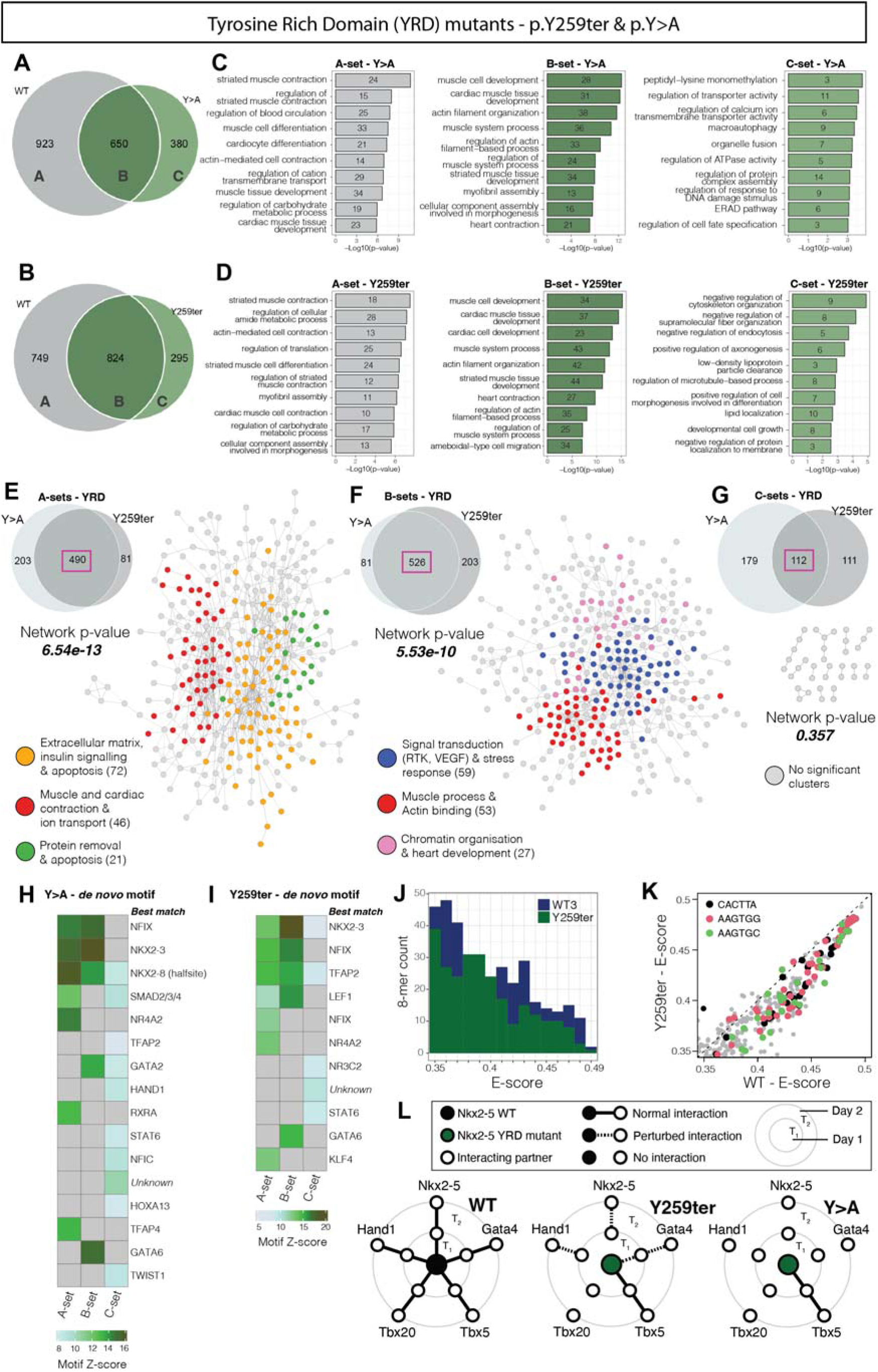
YRD variants result in the broad loss of specific cardiac regulatory programmes due to reduced homo-and hetero-dimerization (A) Overlap of WT and Y>A peaks. (B) Overlap of WT and Y259ter peaks. (C) Top 10 slimmed GO terms for targets of the A-, B-and C-sets from WT vs. Y>A (A). (D) Top 10 slimmed GO terms for targets of the A-, B-and C-sets from WT vs. Y259ter (B). (E) Overlap of genes lost (A-sets) by both YRD variants, from which a high-confidence STRING network was formed with 490 overlapping genes, showing highly significant network p-value, with cardiac network hubs highlighted in red. (F) Overlap of genes retained (B-sets) by both YRD variants, from which a high-confidence STRING network was formed with 526 overlapping genes, showing highly significant network p-value, with cardiac network hubs highlighted in red. (G) Overlap of genes gained (C-sets) by both YRD variants, from which a high-confidence STRING network was formed with 112 overlapping genes, showing no significant network enrichment or key hubs. (H) Significant Trawler de novo enriched motifs clustered to reduce redundancy for A-, B-and C-sets of Y>A variant. (I) Significant Trawler de novo enriched motifs clustered to reduce redundancy for A-, B-and C-sets of Y259ter variant. (J) Counts of significantly bound 8-mers (E > 0.35) from PBM, for WT (dark blue) and Y259ter (green). (K) Sequence specificity of significantly bound 8-mers (E > 0.35) from PBM, for WT against Y259ter, showing no change in specificity with Y259ter. (L) Interaction patterns of Nkx2-5 WT, Y259ter and Y>A variants with cardiac TFs Nkx2-5, Gata4, Tbx5, Tbx20 and Hand1, using a Yeast-two-hybrid (Y2H) protein-protein interaction assay. Y2H fluorescence intensities were normalised to Nkx2-5 WT at 2 different time-points (T_1_ and T_2_) and represented as normal, perturbed or absent interaction plots.

Within the substantial B-set genes for YRD variants (retained targets), we also observed significant enrichment of GO terms for biological processes related to *striated muscle development, differentiation* and *function*, and high overlap, defining a distinct component of the muscle GRN to that detected in A-sets (Figure 3C,D and Figure S7C,D). Interaction analysis of the overlapping B-set genes showed a highly connected network (p-value of 5.53e-10) including a significant core of cardiac-related genes involved in muscle processes, including actin binding (*Ttn*, *Myl1, Des*, *Actc1*), as well as embryonic gene signatures involved in heart development and chromatin organisation (*Tbx20*, *Smarca4*, *Ctcf*) (Figure 3F).

There were also substantial numbers of C-set genes for both YRD variants (295/1119 targets for Y259ter; 380/1030 targets for Y>A) (Figure 3A-B). C-sets represent unique “off-targets” not bound by NKX2-5 WT. Previous work has suggested an underly logic for off-target gene selection by NKX2-5 and other cardiac TF variants using both DamID and ChIP technologies ^25,29^. However, despite high overlap of YRD variant C-set genes studied here, there was little or no network overlap at the GO term level and no cardiac-related terms were evident (Figure 3A-D and S6C-D). Furthermore, STRING network analysis revealed poor interconnectivity of target proteins in the C-sets of Y>A and Y259ter (Table S2), and in the overlap of C-sets (Figure 3G). Together, this suggests that, for YRD variants, the regulatory logic underpinning C-set off-target selection remains hidden.

*De novo* motif discovery was applied to YRD variant A-, B-and C-set target peaks (Figure 2A-B). For both variants, A-set targets (lost targets) showed pronounced overrepresentation of NK2-class (NKE) and NF-1 binding sites, as well as other motifs including those for SMAD2/3/4, NR4A2, TFAP4, RXRA and KLF4 (Figure 3H-I). B-set targets also showed overrepresentation of NKE and NF-1 motifs, suggesting partial disruption of the capacity of variants to bind NKE and NF-1 motifs in core elements of the cardiac network. B-sets uniquely showed overrepresentation of GATA and LEF1 motifs compared to A-sets. C-set targets, which showed no obvious network interconnectivity or logic, nonetheless showed overrepresentation of motifs for TFAP2, STAT6, and HAND1, however, at lower Z-scores. This overall pattern was further corroborated using *de novo* motif analysis tool, Homer ^65^, and known motif enrichment analysis tool, Clover ^96^, which revealed significant differences between A-and B-sets, with fewer motifs in C-sets, confirming a clear regulatory logic for A-and B-sets (Figure S7E-H). In Clover analysis, some NKX and NF-1 motifs were retained in C-sets, but these are likely to be lower affinity. Overall, the Y259ter mutation showed little change in DNA-binding affinity or specificity for high affinity NKE sites compared to NKX2-5 WT, as shown from PBM data (Figure S5D), and in the distribution of all significant 8-mers as a function of E-scores (Figure 3J,K). These analyses powerfully highlight the importance of DNA-binding cofactor interactions in regulatory logic, as seen in the loss and retention of targets in DamID A-and B-sets, respectively, despite normal DNA-binding preference in PBM.

We have previously shown that the YRD has a key role in facilitating NKX2-5 homodimerization (previously thought to occur exclusively through the HD) and heterodimerization, demonstrated for TFs ELK1 and ELK4 ^25^. We therefore systematically profiled the homo-and heterodimerization capabilities of NKX2-5 WT and YRD variants using a quantative liquid-based yeast 2-hybrid (Y2H) protein-protein interaction assay, exploring binding between NKX2-5 WT and cardiac cofactors GATA4, TBX5, TBX20 and HAND1, with data collected at two time points (day 1 and 2) (Figure 3L and S6I). NKX2-5 WT demonstrated sustained or increasing interactions over time with itself and all tested cardiac TFs, known to interact directly with NKX2-5 ^25^. In contrast, both Y259ter and Y>A variants exhibited significantly disrupted or complete loss of interactions with NKX2-5 WT and cardiac co-factors, with the exception of TBX5, which is known to interact through the NKX2-5 HD ^29^ (Figure 3L). These findings suggest that many of the core cardiac TFs interact with NKX2-5 through the YRD. Interestingly, the synthetic Y>A variant, in which all nine tyrosines of the YRD have been substituted for alanine ^79^, showed more profound loss of interaction with NKX2-5 and cofactors compared to the natural Y259ter variant, which impacts only the last of the nine tyrosines. These results again reinforce that cofactor interactions via the YRD play a critical role in the selection of *in vivo* target-sites by NKX2-5 in the presence of an intact HD.

### Homeodomain DNA-backbone-contacting residue variants result in a broad loss of targets

We next focussed on CHD variants affecting DNA backbone phosphate-contacting residues, namely R189G and R190H. Compared to YRD and other HD variants, DNA backbone-contacting variants showed greater severity, demonstrated by greater loss of NKX WT targets (increased A-sets; 82% and 63% of targets, respectively), smaller B-sets (retained targets) and negligible C-sets (Figure 4A-B). As for YRD variants, GO term enrichment analysis using CompGO showed highly significant *muscle* and *actin cytoskeleton*-related signatures in A-and B-set genes (Figure S8A-D). In A-sets, there was also enrichment for terms related to more peripheral aspects of the NKX2-5 regulatory network governing *metabolic* and *catabolic* processes, *protein translation*, *dephosphorylation*, *apoptosis* and *cellular morphogenesis* (Figure 4C-D and S8C-D). Backbone-contacting variant A-sets showed ∼85% intersection, suggesting a shared mechanism for target loss within this functional class (Figure 4E). Generally, overlap of A-set genes within functional classes was much higher than between classes (70-85% versus 35-55%), highlighting that this is network driven. PPI analyses using STRING highlighted less cardiac-related gene nodes in A-sets for backbone-contacting variants compared to YRD variants, and a shift towards loss of more peripheral NKX2-5 network genes involving *RNA metabolism*, *insulin signalling* and *apoptosis* (Figure 4E). B-set target genes were predominantly linked to *muscle development* and *actin filament movement* in both GO term (Figure 4C,D and S8C,D) and network analyses (Figure 4F). Thus, as for YRD variants, backbone-contacting variants maintained an ability to bind core elements of the cardiac GRN, despite broad loss of targets. This was supported by pronounced over-representation of the NK2-class and NF-1 motifs in A-and B-set peaks (Figure 4H,I and S8E-H). Off-target C-set genes of backbone-contacting variants, while overlapping, showed distinct GO terms (Figure 4C,D and S8C,D) which lacked network connectivity (Figure 4G), although this may be influenced by the relatively smaller C-set size (Table S2). Corroborating previous findings using EMSA ^107^, PBM demonstrated that the two backbone contacting variants showed diminished DNA-binding affinity (Figure 4J,K and S5C,D), potentially stemming from compromised structural stability and/or loss of non-specific interactions with DNA ^54,108^. Y2H analysis revealed that R189G and R190H variants showed disrupted interactions with NKX2-5 itself and all cardiac co-factors, save for GATA4 in the case of R189G (Figure 4L-M and S8I). Combined with analysis of YRD variants, these data suggest the profound involvement of both the HD and YRD in NKX2-5 homodimerization and cofactor heterodimerization.

**Figure 4.**
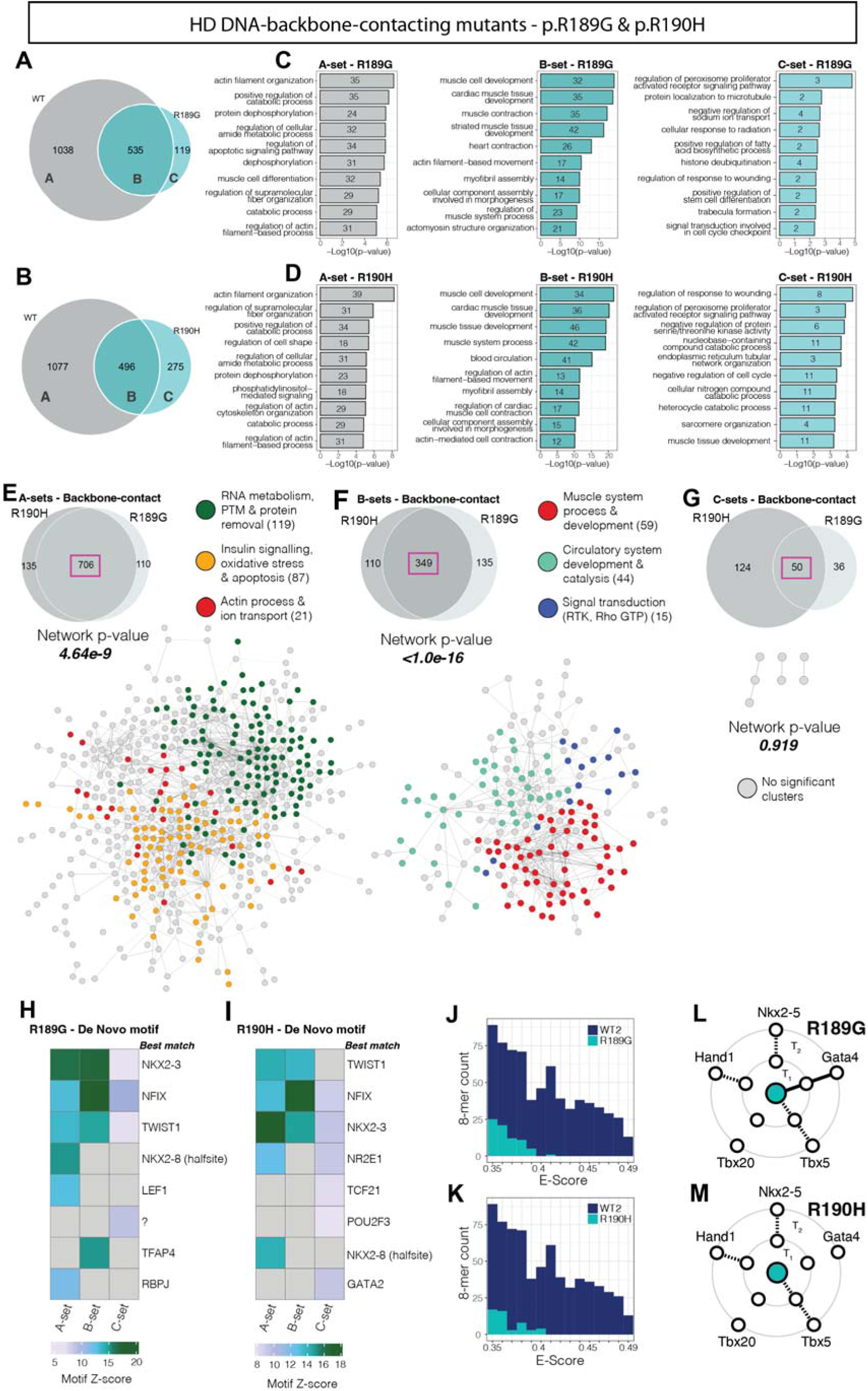
HD DNA-backbone-contacting variants broadly lose many target due to loss of DNA-binding and homo-and hetero-dimerization (A) Overlap of WT and R189G peaks. (B) Overlap of WT and R190H peaks. (C) Top 10 slimmed GO terms for targets of the A-, B-and C-sets from WT vs R189G. (D) Top 10 slimmed GO terms for targets of the A-, B-and C-sets from WT vs. R190H. (E) Overlap of genes lost (A-sets) by both backbone-contacting variants, from which a high-confidence STRING network was formed with 706 overlapping genes, showing highly significant network p-value, with cardiac network hubs highlighted in red. (F) Overlap of genes retained (B-sets) by both backbone-contacting variants, from which a high-confidence STRING network was formed with 349 overlapping genes, showing highly significant network p-value, with cardiac network hubs highlighted in red. (G) Overlap of genes gained (C-sets) by both YRD variants, from which a high-confidence STRING network was formed with 50 overlapping genes, showing no significant network enrichment or key hubs. (H) Significant Trawler de novo enriched motifs clustered to reduce redundancy for A-, B-and C-sets of R189G variant. (I) Significant Trawler de novo enriched motifs clustered to reduce redundancy for A-, B-and C-sets of R190H variant. (J and K) Counts of significantly bound 8-mers (E > 0.35) from PBM, for WT (dark blue) R189G (J) and R190H (K) (light blue). (L and M) Interaction patterns of R189G (L) and R190H (M) variants with cardiac TFs Nkx2-5, Gata4, Tbx5, Tbx20 and Hand1, using a Yeast-two-hybrid (Y2H) protein-protein interaction assay. Y2H fluorescence intensities were normalised to Nkx2-5 WT at 2 different time-points (T_1_ and T_2_) and represented as normal, perturbed or absent interaction plots.

### Homeodomain DNA-base-contacting variants gain cardiac targets through enhanced co-operativity with cardiac TFs and altered DNA binding

All analysed base-contacting residues except R142 were situated within the third alpha helix of the NKX2-5 HD (Figure 1B). There were smaller A-sets for base-contacting variants compared to DNA backbone contacting variants, with N188K exhibiting the greatest loss (Figure 5A-C). Overall, base contacting variant target genes maintained a relatively higher GO term similarity to WT (R=0.69-0.85; Figure S9A-C). Lost targets (A-sets) were associated with *actin filament organisation* terms but predominantly with more peripheral network processes such as *catabolism*, *apoptosis*, *regulation of kinase activity* and *dephosphorylation*, and *translation*, similar to backbone-contacting variant A-sets (Figure S9D-F). The relatively higher number of retained (B-set) genes were consistently associated with *muscle* and *cardiac* GO terms, and these had much higher p-values than for A-set gene terms (Figure S9D-F), the same pattern as observed across other variant functional classes (Figure 3A-D; Figure 5A-D).

**Figure 5.**
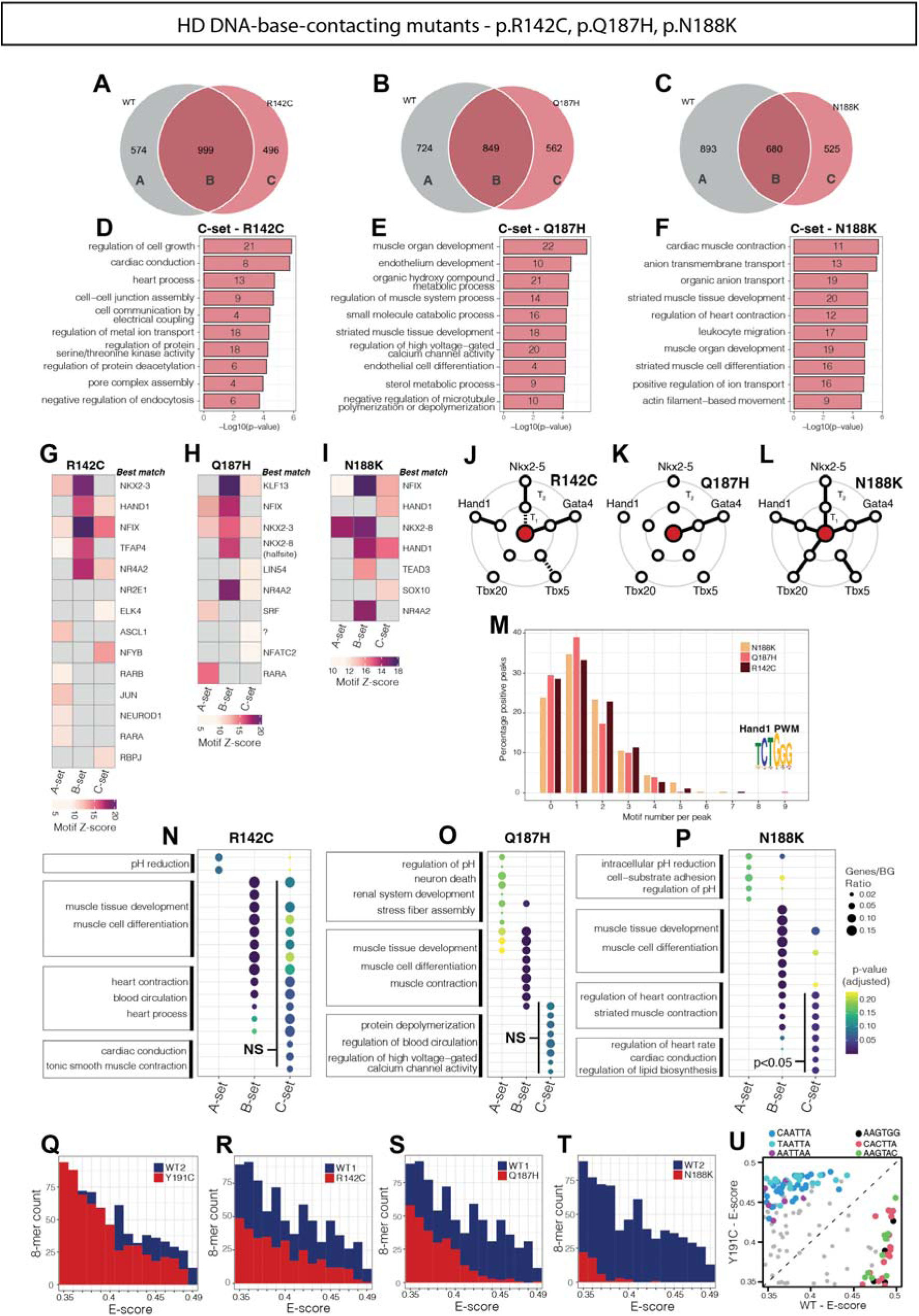
HD DNA-base-contacting variants are redistributed to ectopic cardiac targets via atypical interactions with cardiac TFs (A) Overlap of WT and R142C peaks. (B) Overlap of WT and Q187H peaks. (C) Overlap of WT and N188K peaks. (D) Top 10 slimmed GO terms for targets of the R142C C-set only. (E) Top 10 slimmed GO terms for targets of the Q187H C-set only. (F) Top 10 slimmed GO terms for targets of the N188K C-set only. (G) Significant Trawler de novo enriched motifs clustered to reduce redundancy for A-, B-and C-sets of R142C variant. (H) Significant Trawler de novo enriched motifs clustered to reduce redundancy for A-, B-and C-sets of Q187H variant. (I) Significant Trawler de novo enriched motifs clustered to reduce redundancy for A-, B-and C-sets of N188K variant. (J-L) Interaction patterns of R142C (J), Q187H (K) and N188K (L) variants with cardiac TFs Nkx2-5, Gata4, Tbx5, Tbx20 and Hand1, using a Yeast-two-hybrid (Y2H) protein-protein interaction assay. Y2H fluorescence intensities were normalised to Nkx2-5 WT at 2 different time-points (T_1_ and T_2_) and represented as normal, perturbed or absent interaction plots. (M) Count distribution of Hand1 motifs in C-set peaks of major base-contacting variants. (N-P) GO term analysis of genes associated with high Hand1 motif content peaks (3+ motifs per peak) in base-contacting variant A-, B-and C-sets of R142C (N), Q187H (O) and N188K (P), with C-set terms showing significance only for N188K. (Q-T) Counts of significantly bound 8-mers (E > 0.35) from PBM, for WT (dark blue) and base-contacting variants (red), Y191C (Q), R142C (R), Q187H (S) and N188K (T). (U) Sequence specificity of significantly bound 8-mers (E > 0.35) from PBM, for WT against Y191C, showing a change in specificity with Y191C.

What most distinguished DNA base-contacting variants from other classes was their binding to a higher number of off-targets (C-sets) (Figure 5A-C). C-sets for R142C, Q187H, and N188K represented 33%, 40% and 40% of total targets, respectively. Unlike for other variant classes, genes associated with base-contacting C-set targets were uniquely enriched for a diverse array of cardiomyocyte process, including *muscle development* and *differentiation*, *contraction* and *electrical coupling*, as well as heart-related processes such as *endothelial development* and *metabolism* (Figure 5D-F). TF genes such as *Sox6*, *Nr2f2*, *Smad7*, *Hdac7*, *Hdac9*, *Fos*, *Mef2a*, *Nkx2-5*, *Etv1/2*, as well as various ion channel and intercellular junction genes contributed to the cardiac and endothelial C-set GO terms (Figure S10A-C). N188K was noteworthy in that it bound to targets associated with different membrane transporters, many of which are members of the solute carrier (*Slc*) gene family implicated in transport of organic ions and small molecules including amino acids, neurotransmitters and vitamins ^109^ (Figure S10C). Similar to other variant classes, A-and B-sets of base-contacting variants showed a high degree of overlap, highlighting again the shared mechanisms for target loss and retention in this class (Figure S9H-J). However, despite sharing muscle-related terms, C-set overlap was weaker (as for other functional classes), suggesting that off-target binding generally occurs in a stochastic manner or is highly dependent on the specific residue affected.

Analysis of *de novo* motifs using Trawler revealed that NKE and NF-1 motifs were highly over-represented in B-set peaks of all base-contacting variants (Figure 5G-I), signifying a conserved interaction with essential cardiac regulatory elements. These motifs were less enriched in A-and C-sets. Enrichment analysis of *de novo* motifs with Homer (Figure S11A-C) and known motifs using Clover (Figure S11D-F) provided a similar picture, with the distinction that various NKE sites were enriched in almost all A-sets but absent in C-sets. Focusing on C-set off-target peaks, the most enriched *de novo* discovered motifs were for HAND, NF1, NFY, NR4, NKX2 and KLF TF families (Figure 5G-I), NFY family proteins notable for their function as pioneer TFs ^110^. Known motif analysis in base-contacting C-sets revealed motifs for GATA, SRF, MEF2, TAL1, p53, FOXO, MYB, STAT, AP1, EGR1 and MYOD families, some of which are known cofactors of NKX2-5^25–28,111,112^ (Figure S11A-D).

Y2H analysis for cardiac co-factor interactions illustrated that each of the base-contacting variants presented a highly distinct pattern of interaction disruption (Figure 5J-L and S12). All variants maintained an interaction with GATA4, possibly clarifying the consistent presence of GATA motifs in both Clover and Homer analyses, as well as the enrichment of cardiac targets in variant C-sets. Paradoxically, N188K, the base contacting variant most distinct from NKX2-5 WT, maintained robust interactions with all tested co-factors (weakly for TBX5). Furthermore, this variant showed a particularly strong and persistent interaction with HAND1, which may account for the notable enrichment of HAND1 motifs within its C-set compared to A-set and B-set peaks. We extended the Y2H assay out to day 5, observing a sustained interaction between N188K and HAND1, which for other base-contacting variants ceased after day 2 (Figure S12). Thus, N188K can be characterised as a super-interactor with HAND1.

Total N188K DamID peaks did not display a significant preference for those with higher HAND1 motif content compared with other base-contacting variants (Figure 5M). However, Trawler *de novo* motif enrichment and Clover known motif enrichment highlighted HAND1 motifs significantly enriched in N188K C-set peaks (Figure 5I and S10C,F). We filtered A-, B-and C-sets of base-contacting variants for peaks that contained three or more HAND1 motifs and conducted GO term analysis on associated genes. Whereas filtered A-set genes of base-contacting variants show no significant associations with cardiac GO terms, B-set gene showed a strong association, in-keeping with HAND1 being embedded in the NKX2-5 regulatory network at a high level ^113^ (Figure 5N-P). Among filtered C-set genes, only those of the HAND super-interactor N188K showed significant enrichment (adjusted p-value <0.05). Thus, the super-interaction between N188K and HAND1 provides a mechanistic rationale for the observed N188K gain-of-function.

PBM data revealed that base-contacting variants display varying degrees of loss of DNA-binding affinity to 8-mers from partial (R142C), to almost complete (N188K) (Figure 5Q-T). Consistent with our previous DamID analysis ^25^, PBM analysis of the HD specificity-determining variant, Y191C, affecting one of the three classical specificity-determining residues in helix 3 of the HD, revealed a distinct shift in DNA binding specificity favouring a motif reminiscent of HOX proteins, however, without significant loss of affinity to NKE-containing 8 mers (Figure 5Q and U). These data together reinforce that C-set peaks can be selected via a specific regulatory logic involving atypical cofactor interactions, and/or by changes in DNA binding specificity ^25^.

### Nuclear target search strategy is disrupted by HD DNA-contacting variants

Mammalian TFs spend >50% of their time searching chromatin for their functional DNA targets and their search efficiency is profoundly affected by chromatin structure and cofactor interactions ^54^. To elucidate the chromatin binding and target search functions of NKX2-5 WT and its variants, we implemented a dual-modality single molecule tracking (SMT) imaging approach, similar to that of McCann and colleagues ^71^. This consisted of transient expression of NKX2-5 WT and variants N-terminally fused to the HaloTag self-labelling enzyme in Hela cells^114^. Following addition of the dye JF549, which covalently binds to the expressed HaloTag protein, we imaged single molecules using highly inclined and laminated optical sheet super-resolution (HILO) microscopy ^72,115^. The dual-modality imaging approach included capturing single molecules at two different acquisition rates: “fast” tracking at 20 ms intervals, and “slow” tracking at 500 ms intervals (Figure 6A,B and Figure S13). Fast-tracking broadly capturing both mobile and immobile TF behaviours, and slow-tracking capturing immobile TF persistence (stable binding) ^99^ (Figure S13 and S14). We assessed the nuclear search dynamics of representative examples of the three key functional classes of NKX2-5 variants, namely, base-contacting, backbone-contacting and YRD variants. Previously characterised variant, Y191C, was included as an example of a variant affecting DNA-binding specificity ^25^.

**Figure 6.**
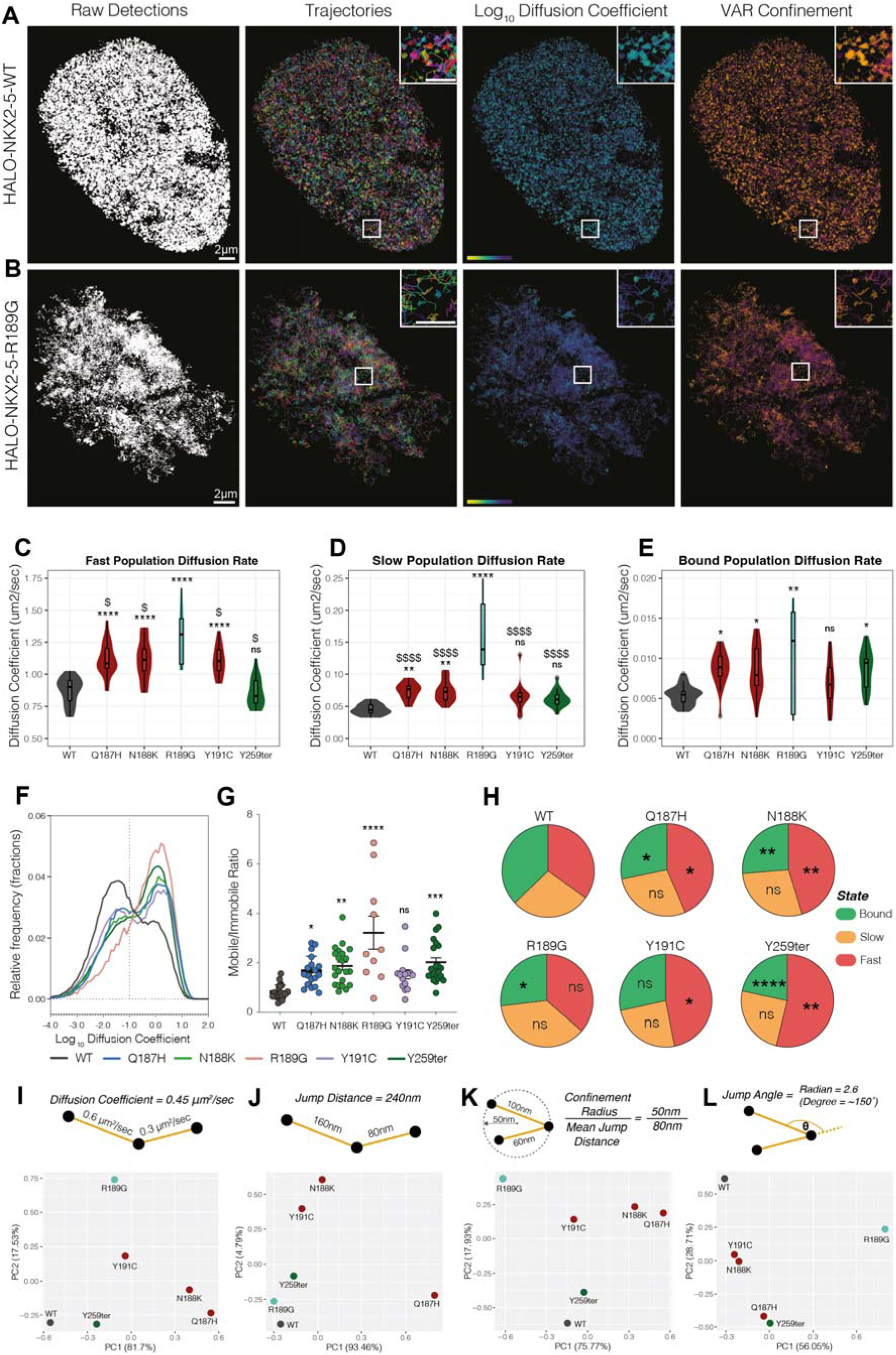
Single molecule tracking reveals divergent dynamics of DNA-base and DNA-backbone contacting NKX2-5 variants (A) Representative analysed super-resolution images of Halo-NKX2-5-WT and (B) Halo-NKX2-5-R189G mutant, showing tracked molecules from a single HeLa nucleus. Panels from left to right: 2D plots of all x,y,t single molecule detections (white), corresponding trajectories (arbitrarily coloured), trajectories colour-coded by log_10_ diffusion coefficient (yellow = lower values/mobilities; purple = higher values/mobilities), and trajectories colour-coded by confinement status as determined by vector autoregression (VAR) analysis (orange = confined; purple = unconfined). Panels in A and B were generated using a custom-made Python software (NASTIC). (C) Diffusion coefficient of fast-diffusing population of NKX2-5 WT or mutant molecules. (D) Diffusion coefficient of slow-diffusing population of NKX2-5 WT or mutant molecules. (E) Diffusion coefficient of stably bound population of NKX2-5 WT or mutant molecules. (F) Distribution of Log_10_ diffusion co-efficient for NKX2-5 WT and mutant molecules, split into mobile and immobile fractions, separated by dotted line. (G) Ratio of NKX2-5 WT or mutant molecules classified as mobile vs. immobile. (H) Percentages of trajectories categorised as either bound, slow-or fast-diffusing for NKX2-5 WT or mutant molecules. (I-L) Principal components analysis of molecular dynamics of NKX2-5 WT or mutant molecules analysed using TrackIt, showing diffusion coefficient per trajectory (I), mean jump distance per trajectory (J), confinement radius over mean jump distance per trajector (K) and jump angle per trajectory (L). Significance (C-E) was calculated with a Two-way ANOVA, * indicates difference from WT less than p<0.05, ** indicates difference from WT less than p<0.01 and **** indicates difference from WT less than p<0.0001. $ indicates difference from backbone-contacting mutants less than p<0.05 and. $$$$ indicates difference from backbone-contacting mutants less than p<0.0001. **Declaration of generative AI and AI-assisted technologies in the writing process.** No generative AI tools were used to write this manuscript.

Analysis of fast tracking data using SpotOn ^77^ allowed segregation of molecules into distinct populations or “states,” which included fast-moving, slow-moving, and bound, and calculated their average diffusion coefficients per cell (Figure 6C-E). Analysis revealed that variant molecules generally exhibited a higher diffusion coefficient than WT, with R189G (severe backbone-contacting variant) the most severely affected. Further analysis using SLIMFast ^75^ generated total log diffusion coefficient curves, capturing the global behaviour of WT and variants (Figure 6F,G). All variants showed deviation from the WT log diffusion coefficient curve, with R189G displaying the most skewed curve and, accordingly, mobile/immobile ratio (Figure 6A-E; Figure S14). This suggests that the R189G and other variants severely disrupt the ability of NKX2-5 to efficiently search the nuclear space for its target sites. Log diffusion coefficient profiles for Y191C and Y259ter, with intact HD folds, were also highly skewed.

Differences in the proportion of molecules occupying each state were determined (Figure 6H) and showed that all variants spent more time in faster moving states and less time strongly bound. Intriguingly, the otherwise severe variant, R189G, was the least severely affected in this assay, whereas Y259ter exhibited the most substantial loss of strong binding state proportion, despite having an intact HD.

We used the slow-tracking imaging modality to evaluate the residence times of NKX2-5 WT and variants in long-and short-lived dwell states, which represent binding to specific and nonspecific sites, respectively ^78^. Dwell times are a function of the TF-DNA dissociation constant ^54^. NKX2-5 WT showed average long-and short-lived dwell times of 5 sec and 0.9 sec, respectively (Figure S15A-B), comparable to other TFs involved in lineage specification ^74^. Most variants exhibited a reduction in residence time by ∼10-30%. Interestingly, the long-to-short lived ratio remained relatively constant, albeit that R189G, one of the most affected variants, showed a significant deviation (Figure S15C). This suggests that the residence times for short-and long-lived molecules are diminished roughly proportionally for most mutations. Cumulative decay function (CDF) analysis suggests that WT and variants have largely comparable turnover of bound molecules as a function of residence time, with only subtle differences (Figure S15D), except for R189G, which displayed a higher number of very long-lived molecules (>20 sec), an apparent gain-of-function (Figure S15D). This is surprising given the lower propensity of DNA backbone-contacting mutants to bind NKX2-5 WT targets (Figure 4A-G).

Principal component analysis (PCA) of total trajectory length data derived from the average of two consecutive windows of both fast-(20 ms) and slow-tracking (500 ms) modalities (i.e. measuring how long a single trajectory lasted across the imaging window), indicated a very similar distribution of variants along the PC1 axis, which accounted for most variation (Figure S15E,F). The backbone-contacting mutation, R189G, showed the greatest deviation from WT. Base-contacting mutations showed similar, although less extreme, behaviours.

We next employed TrackIt ^76^, a tool that generates detailed information on the 2D behaviour of single molecules, providing a comprehensive view, not only of how fast single NKX2-5 WT and variant molecules move, but how they interact spatially within the nuclear environment. PCA for diffusion coefficient probability confirmed that R189G exhibited the greatest deviation from WT, i.e enhanced nuclear mobility (Figure 6I and S16A). Conversely, Y191C and Y259ter, which have intact HDs, displayed the smallest deviation from WT. Base-contacting mutants showed graded differences from NKX2-5 WT.

We also used TrackIt to analyse jump distances, a measure of how far a molecule travels between frames. Unlike for diffusion coefficient (Figure 6I), jump distance PCA analysis showed that R189G clustered closely to WT (Figure 6J), however, it showed a markedly different jump distance distribution strongly skewed towards larger distances (Figure S16B), possibly because its diminished non-specific interactions with DNA provide less constraint ^54,108^. All base-contacting variants and Y259ter only mildly affected jump distance. We also analysed the molecule confinement radius, which reflects the geographic localisation of a molecule in 2D space. WT trajectories were highly confined, typically showing a confinement radius peak at∼100 nm (Figure 6J). In stark contrast, all variants showed profound effects on spatial confinement, as evidenced by PCA (Figure 6J), and all mutants showed a secondary peak of confinement radius at ∼400-500nm, not seen for NKX2-5 WT (Figure S16C). R189G was most severely affected, with most ∼100nm confinements lost. Thus, both backbone-contacting and base-contacting residues are important for achieving highly confined search volumes ^54,74^.

Trajectory jump angles demonstrate the direction in which a molecule travels after each motion, in 2D search space. PCA showed a distinct deviation from NKX2-5 WT for all variants (Figure 6L), with Y191C and N188K clustering tightly together, and Q187H and Y259ter also (Figure 6K). We analysed the anisotropy of trajectories, focusing on connected trajectory segments with jump distances >125nm, representing primarily freely-diffusing, non-bound tracks. We observed that NKX2-5 WT freely-diffusing molecules displayed high anisotropy, peaking at 180° (Figure S17A-F). This indicates that WT molecules are substantially more likely to travel backwards than forwards after each movement. This directional propensity was severely curtailed in variants. The high anisotropy seen for jump distances >125nm was, however, muted for jump trajectories <125nm (Figure S17G-L).

Together, these analyses provide insights into the altered movement, binding behaviour, and spatial dynamics of NKX2-5 variants within the nuclear environment. This offers, for the first time, insights into potentially diverse mechanisms of action of severe cardiac TF mutations in CHD.

## DISCUSSION

Here we undertook a comprehensive functional-epigenomics approach to better understanding the impacts of NKX2-5 CHD mutations on cardiac developmental networks. Our selection of twelve variants allowed us to explore whether disruption of HD residues with distinct structural features represent a common class related to severe loss-of-function (haploinsufficiency ^41^) or show unique epigenomic impacts based on cognate functionalities.

### General features

All variants showed compelling lost (A-set), retained (B-set) and off-target (C-set) binding. The most severe variants (L171P, R189G, R190H, Y25ter, Y>A), showed greater overall loss of targets (increased A-sets, smaller B-and C-sets), compared to variants with milder impacts. Retained B-set targets consistently showed strong over-representation of cardiac muscle-related GO-terms and networks, indicating preservation of variant binding to core elements of the cardiac GRN, irrespective of severity. Significant shifts in the ratio of binding to NKX2-5 motifs of different affinities and specificity (including HOX-like motifs), appeared to be at least partially buffered *in vivo*, presumably though supportive co-factor and chromatin interactions. Lost targets (A-sets) mostly showed muted association with cardiac muscle terms and were more associated with more peripheral aspects of the cardiac GRN. Strikingly, A-and B-set genes individually showed high overlap when comparing variants linked by common structural and functional properties (backbone-contacting, base-contacting, YRD variants), defining these as variant classes and suggesting a novel basis for CHD causation. C-sets showed lower degrees of overlap and less significant p-values, suggesting higher stochasticity of off-target selection (Figure 5A-F).

### N-terminal variants

N-terminal variants K15I and R25C lie respectively within, and adjacent to, the conserved N-terminal TN repressor domain (Figure 1A). Based on peak and GO term analyses, these variants appeared to be mildly affected (Figure 1C,D). Whereas K15I is classified clinically as *likely pathogenic* by ACMG, it shows weakly penetrant phenotypes and may act as a modifier ^37,84^. Interesting, K15I target gene analysis revealed a gain in the significance of muscle GO terms detected (Figure 1C and E), consistent with loss of repressive function.

### HD stabilising variants

HD stabilising variants T178M and L171P showed strikingly different epigenomic signatures. Whereas T178M is classified as *pathogenic* and association with severe CHD in humans (Figure S1B), it segregated closely with NKX2-5 WT, as well other mild variants (Figure 1C-D). Consistently, T178M showed a relatively mild reduction in DNA binding affinities to the NKE (Figure S5D), however, it showed stronger deviation from WT when considering the broader spectrum of target sequences (Figure 2E). L171P, which introduces a kink into HD helix 2 and shows dominant-negative effects experimentally ^92^, had the greatest loss of targets, divergent GO terms and motif signatures of all variants analysed.

### HD DNA backbone-contacting variants

Severe class-specific molecular phenotypes were seen for NKX2-5 DNA backbone-contacting variants R189G and R190H, likely because of their destabilising effects on HD structure and loss of non-specific binding to DNA ^54,108^. Molecular phenotypes for this class were among the most severe, however, in the context of only partial disruption of PPI with cofactors. Consistently, heterozygous mice carrying the R189G mutation show a complex spectrum of Nkx2-5-related cardiac defects not seen in *Nkx2-5* heterozygous null mice ^118^, clearly demonstrating allele-specific anomalies.

### DNA base-contacting variants

Base-contacting variants showed graded degrees of relationship to NKX2-5 WT (Figure 1C; Figure S4A and B). Despite this, A-and B-sets genes for different base-contacting variants showed a high degree of overlap and GO term concordance (Figure 5A-C; Figure S8H-J), thus defining these as an additional HD variant class. R142C is unusual for a base-contacting variant as it lies N-terminal in the HD, interacting specifically with the DNA minor groove. Whereas it is predicted to be pathogenic, it segregated closely to NKX2-5 WT and other mildly affected variants and showed normal binding to the NKE 8-mer (Figure 1C,D; Figure S4A,B; Figure S5D). However, PPI with cardiac co-factor TBX20 was all but lost, and heavily disrupted for NKX2-5, HAND1 and TBX5 with only a strong interaction to GATA4 being retained. R142C also showed reduced E-scores for binding across top 8-mers and a skewing of the proportion of binding to high versus low affinity sites, and HOX-related motifs, revealed by PBM (Figure 2E and F; Figure 5R). R142C also showed a gain in significance of cardiac muscle-related GO terms (Figure 1E), potentially indicating loss of repression. This complex array of features likely underlies its pathogenicity.

Most severe among base-contacting variants was N188K, which was almost as severe as backbone-contacting mutations (Figure 1C and D). However, N188K displaying a distinct pathogenicity profile, having among the most damaging impacts on binding affinity to the NKE and related lower affinity sites among all variants analysed (Figure S5D; Figure 2C-F; Figure 5T). Paradoxically, it retained PPI to NKX2-5 and cofactors with only a partial diminishment for TBX5. Unique to variant classes analysed, N188K C-set off-targets showed compelling over-representation of cardiac *muscle development*, *differentiation*, *contraction*, *actin filament* and *ion transport* terms and networks (Figure 5A-C; S9A-C). We discovered that N188K was in fact a super-interactor with the co-transcription factor HAND1, providing a likely mechanism for this effect. Accordingly, there was increased cardiac muscle GO-terms detected among all N188K gene targets and a highly significant over-representation of HAND motifs among C-set targets associated with genes involved in cardiac contraction (Figure 1E and Figure 5I,P).

### YRD variants

Variants within the conserved YRD represent yet another distinct variant class with severe molecular phenotypes. The YRD is a PPI domain which participates, along with the HD, in NKX2-5 homodimerization and is also a heterodimerisation interface for co-factors ELK1 and ELK4, which are highly embedded within the cardiac network ^25^. Our work here suggests that the YRD is also involved in PPI with GATA4, TBX20 and HAND1, in collaboration with the HD. We have previously shown that the YRD is essential for the function of NKX2-5 in early heart development ^79^. Unique among variants described, YRD variants showed lost (A-set) target gene sets that were highly enriched for GO terms associated with *cardiac muscle development*, *differentiation* and *contraction* (a feature more typical of B-sets). Thus, YRD variants, despite having an intact HD and near-normal binding site affinity using PBM (Figure 3J,K and Figure S5D), failed to bind to a large subset of muscle-associated targets at the core of the cardiac GRN, as well as to more peripheral targets. These data powerfully reinforce the importance of PPI in enhanceosome assembly and regulation of gene expression ^25,29,49^, and the shared responsibility of the HD and the YRD for core NKX2-5 functions.

### Chromatin search and binding behaviours

Mammalian TFs are believed to find their targets on chromatin through *facilitated diffusion* along DNA nano-fibres involving sliding, hopping and intersegmental transfer, nucleosome binding, condensate formation, and stepwise assembly of oligomeric complexes ^54,108^. These functions likely strike a balance between efficient and comprehensive scanning of DNA within diverse and changeable chromatin volumes and exiting them to continue searching elsewhere. It is important to consider that TF amino acid sequences encode not only specificity and affinity parameters for functional DNA-binding and regulation of gene expression, but also chromatin search parameters, which may be profoundly affected by disease variants. Here, we conducted a deep computational analysis of SMT data on NKX2-5 WT and select variants affecting DNA base-contacting (Q187H, N188K), backbone-contacting (R189G), specificity (Y191C) and YRD residues. All variants showed an increase in mean diffusion coefficients within their mobile, slow-moving and bound fractions, the latter two thought to represent non-specific and specific DNA interactions with chromatin, respectively ^54,108^ (Figure 6C,E and G,H). There was also a decrease in mean long-dwell and short-dwell residency times, reflecting an increase in decay rates and overall reduced DNA-binding time (Figure 6G and Figure S15A). Backbone-contacting variant R189G was the most severely affected, with base-contacting variants Q187H and N118K also showing strong deviation from WT. R189G uniquely showed a higher proportion of very long residency times (>20 sec), an apparent gain of function (Figure S15D). Our findings concur with studies on a limited number of other TFs showing that non-specific DNA binding is an important feature of chromatin search functions and alone can lead to long-term residency events ^74^. Non-specific interactions with chromatin, which may involve direct binding to DNA and/or nucleosomes ^74^, are also a dominant feature of TF binding to mitotic chromosomes ^119^. For some TFs, search functions have been found to be modulated by DNA binding specificity residues or PPI determinants ^74,78,120^. Our findings show that HD backbone-contacting, base-contacting and specificity residues, and non-HD PPI residues, all profoundly participate in chromatin search functions by NKX2-5. The specific impacts of single variants and variant classes are rather similar to each other, and no individual variant showed catastrophic collapse. Based on models ^108^, it seems probably that all variants studied here by SMT, even R198G, retain non-specific chromatin scanning capacity to some extent. Overall, our findings suggest that the scanning and specific target binding functions of NKX2-5 profoundly overlap and utilise the full extent of NKX2-5 functionalities, including the interdependent and separate roles of the HD and YRD. Intriguingly, TF binding sites detected using highly sensitive ChIP methods were found to include low affinity, partial or degenerate binding sites ^29,78^, suggesting that such sites may be important for chromatin search functions. All HD and YRD variants show reduced binding to high and low affinity sites, frequently explored a chromatin volume with a large confinement radius (400-500nm), not seen for NKX2-5 WT.

### Summary and perspective

Our findings show highly individualised epigenomic and SMT signatures for each variant studied, which together span a broad range of NKX2-5 functionalities. Each variant shows a unique combination of lost and retained targets, and unique off-targets. Retained B-set binding may result from the net effect of residual chromatin search DNA binding capacities, and cardiac-relevant TF-cofactor interactions, potentially at super-enhancers that regulate core cardiac network genes ^49,121^. Whereas there may be a degree of buffering of the DNA binding affinity or specificity defects associated with variants, as described here, it seems likely that B-set will show abnormal reglulation. Thus, lost A-sets, altered expression of retained B-set target and off-targets all have the potential to destabilise cardiac GRNs, a finding that can be generalised to other disease-causing TF mutations. C-set binding, as demonstrated here and previously ^25^, can be underpinned by changes to DNA binding site specificity and an altered balanced of PPI, whereby mutant TFs can be hijacked to non-cardiac or additional cardiac sites. Luna-Zurita *et al*. found that “ectopic” TF targets were expressed in embryonic stem cell-derived cardiomyocytes mutant for NKX2-5, GATA4 or TBX5, due to the redistribution of respective co-factors across the genome ^29^, potentially analogous to the off-target effects described here. Thus, the optimal stoichiometry of TF, cofactors and chromatin targets prevents binding and activation of ectopic enhancers, a finding with profound implications for how genetic variants cause CHD.

Despite unique epigenomic signatures for each variant, we discerned class-specific epigenetic behaviours related to DNA backbone-contacting, DNA base-contacting and YRD variants, discriminated at single residue resolution. Class-specific epigenomic signatures do not appear to translate cleanly into clinical genotype-phenotype correlations, perhaps masked by the overall impacts of haploinsufficiency, including dominant-negative effects and increased network instability common to severe variants. All variants compromised chromatin search functions, with potential gain-of-functions also detected. Search function defects will likely strongly influence DNA-binding efficiency and chromatin dwell times *in vivo*. Our study reveals that complex loss-of-function, retained function, and gain-of-function effects underpin CHD.

### Limitations

DamID data collected from the mouse atrial HL1 cell line will not capture dynamic target gene changes during development. SMT data was collected in HeLa cells in which TF search and binding behaviours, and chromatin accessibility, may differ from those in cardiac cells. Given complex parameters affecting TF search functions and unique experimental settings, data from different studies may not be directly comparable ^54,108^. DamID and ChIP-related methods sample the genome in different ways and at different stoichiometries, and outcomes may not be directly comparable ^25^. Differences may relate to the different degrees to which each method samples non-specific and weakly-specific targets involved in chromatin search functions^78^. Bioinformatics processes are operator and software-dependent, as described previously ^122^, and results will require confirmation, including in human tissues.

## Acknowledgements

Romaric Bouveret and Ashley van Waardenberg conducted DamID profiling and initial analysis, Matthew Graus helped with SMT data analysis, Yi Zeng with NKX2-5 HD structural predictions and Michael See with GEO submission. This work was supported by the National Health and Medical Research Institute (NHMRC) Australia (1061539, 1074386, 1121172, 2008743, 2007896), Australian Research Council (ARC) (DP0988507; DP210102134), U.S. National Institutes of Health (R01 HG010501). The Novo Nordisk Foundation Center for Stem Cell Medicine is supported by Novo Nordisk Foundation grant NNF21CC0073729. R.P.H. held an NHMRC Australia Fellowship (573705) and NHMRC Senior Principal Research Fellowship (1118576). M.R. held an NHMRC/Australian Heart Foundation Career Development Fellowship (CR 12S 6782) and Australian Heart Foundation Future Leader Fellowship (107328). K.H.K held an A*STAR National Science Scholarship.

## Author contributions

A.W. Conceptualisation, Methodology, Validation, Formal Analysis, Data Curation, Writing – Original Draft Preparation, Writing – Review and Editing, Visualisation, Supervision, Project Administration.

N.S. Methodology, Validation, Investigation, Writing – Review and Editing.

A.J.M. Methodology, Formal Analysis, Investigation, Writing – Review and Editing.

S.K.P. Investigation.

K.H.K. Investigation.

J.V.K. Investigation.

S.M. Investigation.

F.W. Investigation.

N.M. Investigation.

J.W. Investigation.

D.A. Formal Analysis, Visualisation.

F.A.M. Resources, Supervision, Writing – Review and Editing.

M.F. Resources, Supervision, Writing – Review and Editing.

M.L.B. Formal Analysis, Resources, Writing – Review and Editing, Supervision, Project Administration, Funding Acquisition.

M.R. Formal Analysis, Supervision, Writing - Review and Editing.

R.P.H. Conceptualisation, Methodology, Resources, Writing – Original Draft Preparation, Writing – Review and Editing, Supervision, Project Administration, Funding Acquisition.

## Data Availability

Raw data has been deposited at the NCBI Gene Expression Omnibus (GEO) database under accession number: GSE318589

## Competing interests

M.L.B. is a co-inventor on U.S. patents #6,548,021 and #8,530,638 on PBM technology and corresponding universal sequence designs, respectively. Universal PBM array designs used in this study are available via a Materials Transfer Agreement with The Brigham & Women’s Hospital, Inc. The remaining authors declare no competing interests.

## Notes

https://www.ncbi.nlm.nih.gov/geo/query/acc.cgi?&acc=GSE318589

### Competing Interest Statement

M.L.B. is a co-inventor on U.S. patents 6548021 and 8530638 on PBM technology and corresponding universal sequence designs, respectively. Universal PBM array designs used in this study are available via a Materials Transfer Agreement with The Brigham & Women's Hospital Inc. The remaining authors declare no competing interests.

### Summary of Updates

Text substantially revised and figures updated.

## References

1 Kirk, E. P. et al. Mutations in cardiac T-box factor gene TBX20 are associated with diverse cardiac pathologies, including defects of septation and valvulogenesis and cardiomyopathy. Am J Hum Genet 81, 280–291 (2007). 10.1086/519530

2 Schott, J. J. et al. Congenital heart disease caused by mutations in the transcription factor NKX2-5. Science 281, 108–111 (1998). 10.1126/science.281.5373.108

3 Basson, C. T. et al. Mutations in human TBX5 [corrected] cause limb and cardiac malformation in Holt-Oram syndrome. Nat Genet 15, 30–35 (1997). 10.1038/ng0197-30

4 Li, Q. Y. et al. Holt-Oram syndrome is caused by mutations in TBX5, a member of the Brachyury (T) gene family. Nat Genet 15, 21–29 (1997). 10.1038/ng0197-21

5 Garg, V. et al. GATA4 mutations cause human congenital heart defects and reveal an interaction with TBX5. Nature 424, 443–447 (2003). 10.1038/nature01827

6 Chapman, G. et al. Functional genomics and gene-environment interaction highlight the complexity of congenital heart disease caused by Notch pathway variants. Hum Mol Genet 29, 566–579 (2020). 10.1093/hmg/ddz270

7 Barnes, R. M. & Black, B. L. in Etiology and Morphogenesis of Congenital Heart Disease: From Gene Function and Cellular Interaction to Morphology (eds T. Nakanishi et al.) 183–192 (2016).

8 Tan, H. L. et al. Nonsynonymous variants in the SMAD6 gene predispose to congenital cardiovascular malformation. Hum Mutat 33, 720–727 (2012). 10.1002/humu.22030

9 Klena, N. T., Gibbs, B. C. & Lo, C. W. Cilia and Ciliopathies in Congenital Heart Disease. Cold Spring Harb Perspect Biol 9 (2017). 10.1101/cshperspect.a028266

10 Hilal, N., Chen, Z., Chen, M. H. & Choudhury, S. RASopathies and cardiac manifestations. Front Cardiovasc Med 10, 1176828 (2023). 10.3389/fcvm.2023.1176828

11 Zaidi, S. et al. De novo mutations in histone-modifying genes in congenital heart disease. Nature 498, 220–223 (2013). 10.1038/nature12141

12 Homsy, J. et al. De novo mutations in congenital heart disease with neurodevelopmental and other congenital anomalies. Science 350, 1262–1266 (2015). 10.1126/science.aac9396

13 Jin, S. C. et al. Contribution of rare inherited and de novo variants in 2,871 congenital heart disease probands. Nat Genet 49, 1593–1601 (2017). 10.1038/ng.3970

14 Sifrim, A. et al. Distinct genetic architectures for syndromic and nonsyndromic congenital heart defects identified by exome sequencing. Nat Genet 48, 1060–1065 (2016). 10.1038/ng.3627

15 Ng, S. B. et al. Exome sequencing identifies MLL2 mutations as a cause of Kabuki syndrome. Nat Genet 42, 790–793 (2010). 10.1038/ng.646

16 Vissers, L. E. et al. Mutations in a new member of the chromodomain gene family cause CHARGE syndrome. Nat Genet 36, 955–957 (2004). 10.1038/ng1407

17 Robson, A. et al. Histone H2B monoubiquitination regulates heart development via epigenetic control of cilia motility. Proc Natl Acad Sci U S A 116, 14049–14054 (2019). 10.1073/pnas.1808341116

18 Pierpont, M. E. et al. Genetic Basis for Congenital Heart Disease: Revisited: A Scientific Statement From the American Heart Association. Circulation 138, e653–e711 (2018). 10.1161/CIR.0000000000000606

19 Davidson, E. H. & Erwin, D. H. Gene regulatory networks and the evolution of animal body plans. Science 311, 796–800 (2006). 10.1126/science.1113832

20 Nagy, G. & Nagy, L. Motif grammar: The basis of the language of gene expression. Comput Struct Biotechnol J 18, 2026–2032 (2020). 10.1016/j.csbj.2020.07.007

21 Buecker, C. & Wysocka, J. Enhancers as information integration hubs in development: lessons from genomics. Trends Genet 28, 276–284 (2012). 10.1016/j.tig.2012.02.008

22 Hoppe, C. et al. Modulation of the Promoter Activation Rate Dictates the Transcriptional Response to Graded BMP Signaling Levels in the Drosophila Embryo. Dev Cell 54, 727–741 e727 (2020). 10.1016/j.devcel.2020.07.007

23 Maytum, A., Edginton-White, B. & Bonifer, C. Identification and characterization of enhancer elements controlling cell type-specific and signalling dependent chromatin programming during hematopoietic development. Stem Cell Investig 10, 14 (2023). 10.21037/sci-2023-011

24 O’tsuki, L., Plattner, S. A., Taniguchi-Sugiura, Y., Falcon, F. & Tanaka, E. M. Molecular basis of positional memory in limb regeneration. Nature (2025). 10.1038/s41586-025-09036-5

25 Bouveret, R. et al. NKX2-5 mutations causative for congenital heart disease retain functionality and are directed to hundreds of targets. Elife 4 (2015). 10.7554/eLife.06942

26 Waardenberg, A. J., Homan, B., Mohamed, S., Harvey, R. P. & Bouveret, R. Prediction and validation of protein-protein interactors from genome-wide DNA-binding data using a knowledge-based machine-learning approach. Open Biol 6 (2016). 10.1098/rsob.160183

27 He, A., Kong, S. W., Ma, Q. & Pu, W. T. Co-occupancy by multiple cardiac transcription factors identifies transcriptional enhancers active in heart. Proc Natl Acad Sci U S A 108, 5632–5637 (2011). 10.1073/pnas.1016959108

28 Akerberg, B. N. et al. A reference map of murine cardiac transcription factor chromatin occupancy identifies dynamic and conserved enhancers. Nat Commun 10, 4907 (2019). 10.1038/s41467-019-12812-3

29 Luna-Zurita, L. et al. Complex Interdependence Regulates Heterotypic Transcription Factor Distribution and Coordinates Cardiogenesis. Cell 164, 999–1014 (2016). 10.1016/j.cell.2016.01.004

30 Lyons, I. et al. Myogenic and morphogenetic defects in the heart tubes of murine embryos lacking the homeo box gene Nkx2-5. Genes Dev 9, 1654–1666 (1995). 10.1101/gad.9.13.1654

31 Prall, O. W. et al. An Nkx2-5/Bmp2/Smad1 negative feedback loop controls heart progenitor specification and proliferation. Cell 128, 947–959 (2007). 10.1016/j.cell.2007.01.042

32 Paffett-Lugassy, N. et al. Heart field origin of great vessel precursors relies on nkx2.5-mediated vasculogenesis. Nat Cell Biol 15, 1362–1369 (2013). 10.1038/ncb2862

33 Pashmforoush, M. et al. Nkx2-5 pathways and congenital heart disease; loss of ventricular myocyte lineage specification leads to progressive cardiomyopathy and complete heart block. Cell 117, 373–386 (2004). 10.1016/s0092-8674(04)00405-2

34 Mommersteeg, M. T. et al. Molecular pathway for the localized formation of the sinoatrial node. Circ Res 100, 354–362 (2007). 10.1161/01.RES.0000258019.74591.b3

35 Jay, P. Y. et al. Nkx2-5 mutation causes anatomic hypoplasia of the cardiac conduction system. J Clin Invest 113, 1130–1137 (2004). 10.1172/JCI19846

36 Karczewski, K. J. et al. The mutational constraint spectrum quantified from variation in 141,456 humans. Nature 581, 434–443 (2020). 10.1038/s41586-020-2308-7

37 Kolomenski, J. E. et al. An update on genetic variants of the NKX2-5. Hum Mutat 41, 1187–1208 (2020). 10.1002/humu.24030

38 Elliott, D. A. et al. Cardiac homeobox gene NKX2-5 mutations and congenital heart disease: associations with atrial septal defect and hypoplastic left heart syndrome. J Am Coll Cardiol 41, 2072–2076 (2003). 10.1016/s0735-1097(03)00420-0

39 Steimle, J. D. & Moskowitz, I. P. TBX5: A Key Regulator of Heart Development. Curr Top Dev Biol 122, 195–221 (2017). 10.1016/bs.ctdb.2016.08.008

40 Whitcomb, J., Gharibeh, L. & Nemer, M. From embryogenesis to adulthood: Critical role for GATA factors in heart development and function. IUBMB Life 72, 53–67 (2020). 10.1002/iub.2163

41 Johnson, A. F., Nguyen, H. T. & Veitia, R. A. Causes and effects of haploinsufficiency. Biol Rev Camb Philos Soc 94, 1774–1785 (2019). 10.1111/brv.12527

42 Gifford, C. A. et al. Oligogenic inheritance of a human heart disease involving a genetic modifier. Science 364, 865–870 (2019). 10.1126/science.aat5056

43 Liu, X. et al. The complex genetics of hypoplastic left heart syndrome. Nat Genet 49, 1152–1159 (2017). 10.1038/ng.3870

44 Dong, P. & Liu, Z. Shaping development by stochasticity and dynamics in gene regulation. Open Biol 7 (2017). 10.1098/rsob.170030

45 Gillinder, K. R. et al. Promiscuous DNA-binding of a mutant zinc finger protein corrupts the transcriptome and diminishes cell viability. Nucleic Acids Res 45, 1130–1143 (2017). 10.1093/nar/gkw1014

46 Ibrahim, D. M. et al. Distinct global shifts in genomic binding profiles of limb malformation-associated HOXD13 mutations. Genome Res 23, 2091–2102 (2013). 10.1101/gr.157610.113

47 Ilsley, M. D. et al. Corrupted DNA-binding specificity and ectopic transcription underpin dominant neomorphic mutations in KLF/SP transcription factors. BMC Genomics 20, 417 (2019). 10.1186/s12864-019-5805-z

48 Barrera, L. A. et al. Survey of variation in human transcription factors reveals prevalent DNA binding changes. Science 351, 1450–1454 (2016). 10.1126/science.aad2257

49 Ang, Y. S. et al. Disease Model of GATA4 Mutation Reveals Transcription Factor Cooperativity in Human Cardiogenesis. Cell 167, 1734–1749 e1722 (2016). 10.1016/j.cell.2016.11.033

50 Freed-Pastor, W. A. & Prives, C. Mutant p53: one name, many proteins. Genes Dev 26, 1268–1286 (2012). 10.1101/gad.190678.112

51 Zhu, J. et al. Gain-of-function p53 mutants co-opt chromatin pathways to drive cancer growth. Nature 525, 206–211 (2015). 10.1038/nature15251

52 Bailey, C. G. et al. Structure-function relationships explain CTCF zinc finger mutation phenotypes in cancer. Cell Mol Life Sci 78, 7519–7536 (2021). 10.1007/s00018-021-03946-z

53 Kock, K. H. et al. DNA binding analysis of rare variants in homeodomains reveals homeodomain specificity-determining residues. Nat Commun 15, 3110 (2024). 10.1038/s41467-024-47396-0

54 Suter, D. M. Transcription Factors and DNA Play Hide and Seek. Trends Cell Biol 30, 491–500 (2020). 10.1016/j.tcb.2020.03.003

55 Rentzsch, P., Witten, D., Cooper, G. M., Shendure, J. & Kircher, M. CADD: predicting the deleteriousness of variants throughout the human genome. Nucleic Acids Res 47, D886–D894 (2019). 10.1093/nar/gky1016

56 Feng, B. J. PERCH: A Unified Framework for Disease Gene Prioritization. Hum Mutat 38, 243–251 (2017). 10.1002/humu.23158

57 Ioannidis, N. M. et al. REVEL: An Ensemble Method for Predicting the Pathogenicity of Rare Missense Variants. Am J Hum Genet 99, 877–885 (2016). 10.1016/j.ajhg.2016.08.016

58 Vogel, M. J. et al. Human heterochromatin proteins form large domains containing KRAB-ZNF genes. Genome Res 16, 1493–1504 (2006). 10.1101/gr.5391806

59 Vogel, M. J., Peric-Hupkes, D. & van Steensel, B. Detection of in vivo protein-DNA interactions using DamID in mammalian cells. Nat Protoc 2, 1467–1478 (2007). 10.1038/nprot.2007.148

60 Claycomb, W. C. et al. HL-1 cells: a cardiac muscle cell line that contracts and retains phenotypic characteristics of the adult cardiomyocyte. Proc Natl Acad Sci U S A 95, 2979–2984 (1998). 10.1073/pnas.95.6.2979

61 Gu, Z., Eils, R. & Schlesner, M. Complex heatmaps reveal patterns and correlations in multidimensional genomic data. Bioinformatics 32, 2847–2849 (2016). 10.1093/bioinformatics/btw313

62 Waardenberg, A. J., Basset, S. D., Bouveret, R. & Harvey, R. P. CompGO: an R package for comparing and visualizing Gene Ontology enrichment differences between DNA binding experiments. BMC Bioinformatics 16, 275 (2015). 10.1186/s12859-015-0701-2

63 Haudry, Y., Ramialison, M., Paten, B., Wittbrodt, J. & Ettwiller, L. Using Trawler_standalone to discover overrepresented motifs in DNA and RNA sequences derived from various experiments including chromatin immunoprecipitation. Nat Protoc 5, 323–334 (2010). 10.1038/nprot.2009.158

64 Castro-Mondragon, J. A., Jaeger, S., Thieffry, D., Thomas-Chollier, M. & van Helden, J. RSAT matrix-clustering: dynamic exploration and redundancy reduction of transcription factor binding motif collections. Nucleic Acids Res 45, e119 (2017). 10.1093/nar/gkx314

65 Heinz, S. et al. Simple combinations of lineage-determining transcription factors prime cis-regulatory elements required for macrophage and B cell identities. Mol Cell 38, 576–589 (2010). 10.1016/j.molcel.2010.05.004

66 Gupta, S., Stamatoyannopoulos, J. A., Bailey, T. L. & Noble, W. S. Quantifying similarity between motifs. Genome Biol 8, R24 (2007). 10.1186/gb-2007-8-2-r24

67 Szklarczyk, D. et al. The STRING database in 2021: customizable protein-protein networks, and functional characterization of user-uploaded gene/measurement sets. Nucleic Acids Res 49, D605–D612 (2021). 10.1093/nar/gkaa1074

68 Shannon, P. et al. Cytoscape: a software environment for integrated models of biomolecular interaction networks. Genome Res 13, 2498–2504 (2003). 10.1101/gr.1239303

69 Berger, M. F. et al. Compact, universal DNA microarrays to comprehensively determine transcription-factor binding site specificities. Nat Biotechnol 24, 1429–1435 (2006). 10.1038/nbt1246

70 Berger, M. F. & Bulyk, M. L. Universal protein-binding microarrays for the comprehensive characterization of the DNA-binding specificities of transcription factors. Nat Protoc 4, 393–411 (2009). 10.1038/nprot.2008.195

71 McCann, A. J. et al. A dominant-negative SOX18 mutant disrupts multiple regulatory layers essential to transcription factor activity. Nucleic Acids Res 49, 10931–10955 (2021). 10.1093/nar/gkab820

72 Tokunaga, M., Imamoto, N. & Sakata-Sogawa, K. Highly inclined thin illumination enables clear single-molecule imaging in cells. Nat Methods 5, 159–161 (2008). 10.1038/nmeth1171

73 Cheng, M. et al. Single-molecule dynamics of site-specific labeled transforming growth factor type II receptors on living cells. Chem Commun (Camb) 50, 14724–14727 (2014). 10.1039/c4cc02804j

74 Lerner, J. et al. Two-Parameter Mobility Assessments Discriminate Diverse Regulatory Factor Behaviors in Chromatin. Mol Cell 79, 677–688 e676 (2020). 10.1016/j.molcel.2020.05.036

75 Serge, A., Bertaux, N., Rigneault, H. & Marguet, D. Dynamic multiple-target tracing to probe spatiotemporal cartography of cell membranes. Nat Methods 5, 687–694 (2008). 10.1038/nmeth.1233

76 Kuhn, T., Hettich, J., Davtyan, R. & Gebhardt, J. C. M. Single molecule tracking and analysis framework including theory-predicted parameter settings. Sci Rep 11, 9465 (2021). 10.1038/s41598-021-88802-7

77 Hansen, A. S. et al. Robust model-based analysis of single-particle tracking experiments with Spot-On. Elife 7 (2018). 10.7554/eLife.33125

78 Chen, J. et al. Single-molecule dynamics of enhanceosome assembly in embryonic stem cells. Cell 156, 1274–1285 (2014). 10.1016/j.cell.2014.01.062

79 Elliott, D. A. et al. A tyrosine-rich domain within homeodomain transcription factor Nkx2-5 is an essential element in the early cardiac transcriptional regulatory machinery. Development 133, 1311–1322 (2006). 10.1242/dev.02305

80 Harvey, R. P. NK-2 homeobox genes and heart development. Dev Biol 178, 203–216 (1996). 10.1006/dbio.1996.0212

81 Kircher, M. et al. A general framework for estimating the relative pathogenicity of human genetic variants. Nat Genet 46, 310–315 (2014). 10.1038/ng.2892

82 Pradhan, L. et al. Crystal structure of the human NKX2.5 homeodomain in complex with DNA target. Biochemistry 51, 6312–6319 (2012). 10.1021/bi300849c

83 Richards, S. et al. Standards and guidelines for the interpretation of sequence variants: a joint consensus recommendation of the American College of Medical Genetics and Genomics and the Association for Molecular Pathology. Genet Med 17, 405–424 (2015). 10.1038/gim.2015.30

84 McElhinney, D. B., Geiger, E., Blinder, J., Benson, D. W. & Goldmuntz, E. NKX2.5 mutations in patients with congenital heart disease. J Am Coll Cardiol 42, 1650–1655 (2003). 10.1016/j.jacc.2003.05.004

85 van Steensel, B. & Henikoff, S. Identification of in vivo DNA targets of chromatin proteins using tethered dam methyltransferase. Nat Biotechnol 18, 424–428 (2000). 10.1038/74487

86 Cheetham, S. W. et al. Targeted DamID reveals differential binding of mammalian pluripotency factors. Development 145 (2018). 10.1242/dev.170209

87 Aughey, G. N., Cheetham, S. W. & Southall, T. D. DamID as a versatile tool for understanding gene regulation. Development 146 (2019). 10.1242/dev.173666

88 Waldminghaus, T. & Skarstad, K. ChIP on Chip: surprising results are often artifacts. BMC Genomics 11, 414 (2010). 10.1186/1471-2164-11-414

89 Ramialison, M. et al. Analysis of steric effects in DamID profiling of transcription factor target genes. Genomics 109, 75–82 (2017). 10.1016/j.ygeno.2017.01.006

90 Friedman, C. E. et al. HOPX-associated molecular programs control cardiomyocyte cell states underpinning cardiac structure and function. Dev Cell 59, 91–107 e106 (2024). 10.1016/j.devcel.2023.11.012

91 Favorov, A. et al. Exploring massive, genome scale datasets with the GenometriCorr package. PLoS Comput Biol 8, e1002529 (2012). 10.1371/journal.pcbi.1002529

92 Grow, M. W. & Krieg, P. A. Tinman function is essential for vertebrate heart development: elimination of cardiac differentiation by dominant inhibitory mutants of the tinman-related genes, XNkx2-3 and XNkx2-5. Dev Biol 204, 187–196 (1998). 10.1006/dbio.1998.9080

93 McLean, C. Y. et al. GREAT improves functional interpretation of cis-regulatory regions. Nat Biotechnol 28, 495–501 (2010). 10.1038/nbt.1630

94 Ettwiller, L., Paten, B., Ramialison, M., Birney, E. & Wittbrodt, J. Trawler: de novo regulatory motif discovery pipeline for chromatin immunoprecipitation. Nat Methods 4, 563–565 (2007). 10.1038/nmeth1061

95 Dang, L. T. et al. TrawlerWeb: an online de novo motif discovery tool for next-generation sequencing datasets. BMC Genomics 19, 238 (2018). 10.1186/s12864-018-4630-0

96 Frith, M. C. et al. Detection of functional DNA motifs via statistical over-representation. Nucleic Acids Res 32, 1372–1381 (2004). 10.1093/nar/gkh299

97 Berger, M. F. et al. Variation in homeodomain DNA binding revealed by high-resolution analysis of sequence preferences. Cell 133, 1266–1276 (2008). 10.1016/j.cell.2008.05.024

98 Mukherjee, S. et al. Rapid analysis of the DNA-binding specificities of transcription factors with DNA microarrays. Nat Genet 36, 1331–1339 (2004). 10.1038/ng1473

99 Chen, C. Y. & Schwartz, R. J. Identification of novel DNA binding targets and regulatory domains of a murine tinman homeodomain factor, nkx-2.5. J Biol Chem 270, 15628–15633 (1995). 10.1074/jbc.270.26.15628

100 Tell, G. et al. Comparative stability analysis of the thyroid transcription factor 1 and Antennapedia homeodomains: evidence for residue 54 in controlling the structural stability of the recognition helix. Int J Biochem Cell Biol 31, 1339–1353 (1999). 10.1016/s1357-2725(99)00047-3

101 Gordan, R. et al. Genomic regions flanking E-box binding sites influence DNA binding specificity of bHLH transcription factors through DNA shape. Cell Rep 3, 1093–1104 (2013). 10.1016/j.celrep.2013.03.014

102 Dror, I., Rohs, R. & Mandel-Gutfreund, Y. How motif environment influences transcription factor search dynamics: Finding a needle in a haystack. Bioessays 38, 605–612 (2016). 10.1002/bies.201600005

103 Yella, V. R. et al. Flexibility and structure of flanking DNA impact transcription factor affinity for its core motif. Nucleic Acids Res 46, 11883–11897 (2018). 10.1093/nar/gky1057

104 Kim, S. & Wysocka, J. Deciphering the multi-scale, quantitative cis-regulatory code. Mol Cell 83, 373–392 (2023). 10.1016/j.molcel.2022.12.032

105 Szklarczyk, D. et al. STRING v11: protein-protein association networks with increased coverage, supporting functional discovery in genome-wide experimental datasets. Nucleic Acids Res 47, D607–D613 (2019). 10.1093/nar/gky1131

106 van Dongen, S. Graph clustering via a discrete uncoupling process. SIAM Journal on Matrix Analysis and Applications 30, 121–141 (2008). 10.1137/040608635

107 Kasahara, H. et al. Loss of function and inhibitory effects of human CSX/NKX2.5 homeoprotein mutations associated with congenital heart disease. J Clin Invest 106, 299–308 (2000). 10.1172/JCI9860

108 Mazzocca, M., Fillot, T., Loffreda, A., Gnani, D. & Mazza, D. The needle and the haystack: single molecule tracking to probe the transcription factor search in eukaryotes. Biochem Soc Trans 49, 1121–1132 (2021). 10.1042/BST20200709

109 Liu, X. SLC Family Transporters. Adv Exp Med Biol 1141, 101–202 (2019). 10.1007/978-981-13-7647-4_3

110 Oldfield, A. J. et al. Histone-fold domain protein NF-Y promotes chromatin accessibility for cell type-specific master transcription factors. Mol Cell 55, 708–722 (2014). 10.1016/j.molcel.2014.07.005

111 Chen, C. Y. et al. Activation of the cardiac alpha-actin promoter depends upon serum response factor, Tinman homologue, Nkx-2.5, and intact serum response elements. *Dev Genet* 19, 119-130 (1996). 10.1002/(SICI)1520-6408(1996)19:2<119::AID-DVG3>3.0.CO;2-C

112 Durocher, D., Charron, F., Warren, R., Schwartz, R. J. & Nemer, M. The cardiac transcription factors Nkx2-5 and GATA-4 are mutual cofactors. EMBO J 16, 5687–5696 (1997). 10.1093/emboj/16.18.5687

113 Buckingham, M., Meilhac, S. & Zaffran, S. Building the mammalian heart from two sources of myocardial cells. Nat Rev Genet 6, 826–835 (2005). 10.1038/nrg1710

114 Los, G. V. et al. HaloTag: a novel protein labeling technology for cell imaging and protein analysis. ACS Chem Biol 3, 373–382 (2008). 10.1021/cb800025k

115 Grimm, J. B. et al. A general method to improve fluorophores for live-cell and single-molecule microscopy. Nat Methods 12, 244–250, 243 p following 250 (2015). 10.1038/nmeth.3256

116 Wallis, T. P. et al. Super-resolved trajectory-derived nanoclustering analysis using spatiotemporal indexing. Nat Commun 14, 3353 (2023). 10.1038/s41467-023-38866-y

117 Benson, D. W. et al. Mutations in the cardiac transcription factor NKX2.5 affect diverse cardiac developmental pathways. J Clin Invest 104, 1567–1573 (1999). 10.1172/JCI8154

118 Ashraf, H. et al. A mouse model of human congenital heart disease: high incidence of diverse cardiac anomalies and ventricular noncompaction produced by heterozygous Nkx2-5 homeodomain missense mutation. Circ Cardiovasc Genet 7, 423–433 (2014). 10.1161/CIRCGENETICS.113.000281

119 Raccaud, M. et al. Mitotic chromosome binding predicts transcription factor properties in interphase. Nat Commun 10, 487 (2019). 10.1038/s41467-019-08417-5

120 Wagh, K. et al. Dynamic switching of transcriptional regulators between two distinct low-mobility chromatin states. Sci Adv 9, eade1122 (2023). 10.1126/sciadv.ade1122

121 Hnisz, D., Shrinivas, K., Young, R. A., Chakraborty, A. K. & Sharp, P. A. A Phase Separation Model for Transcriptional Control. Cell 169, 13–23 (2017). 10.1016/j.cell.2017.02.007

122 Miranda, A. M. A. et al. Single-cell transcriptomics for the assessment of cardiac disease. Nat Rev Cardiol 20, 289–308 (2023). 10.1038/s41569-022-00805-7

